# An accurate probabilistic step finder for time-series analysis

**DOI:** 10.1101/2023.09.19.558535

**Authors:** Alex Rojewski, Maxwell Schweiger, Ioannis Sgouralis, Matthew Comstock, Steve Pressé

## Abstract

Noisy time-series data is commonly collected from sources including Förster Resonance Energy Transfer experiments, patch clamp and force spectroscopy setups, among many others. Two of the most common paradigms for the detection of discrete transitions in such time-series data include: hidden Markov models (HMMs) and step-finding algorithms. HMMs, including their extensions to infinite state-spaces, inherently assume in analysis that holding times in discrete states visited are geometrically–or, loosely speaking in common language, exponentially–distributed. Thus the determination of step locations, especially in sparse and noisy data, is biased by HMMs toward identifying steps resulting in geometric holding times. In contrast, existing step-finding algorithms, while free of this restraint, often rely on *ad hoc* metrics to penalize steps recovered in time traces (by using various information criteria) and otherwise rely on approximate greedy algorithms to identify putative global optima. Here, instead, we devise a robust and general probabilistic (Bayesian) step-finding tool that neither relies on *ad hoc* metrics to penalize step numbers nor assumes geometric holding times in each state. As the number of steps themselves in a time-series are, *a priori* unknown, we treat these within a Bayesian nonparametric (BNP) paradigm. We find that the method developed, Bayesian Nonparametric Step (BNP-Step), accurately determines the number and location of transitions between discrete states without any assumed kinetic model and learns the emission distribution characteristic of each state. In doing so, we verify that BNP-Step can analyze sparser data sets containing higher noise and more closely-spaced states than otherwise resolved by current state-of-the-art methods. What is more, BNP-Step rigorously propagates measurement uncertainty into uncertainty over state transition locations, numbers, and emission levels as characterized by the posterior. We demonstrate the performance of BNP-Step on both synthetic data as well as data drawn from force spectroscopy experiments.

**SIGNIFICANCE:** Many time-series data sets exist which are challenging to analyze with current state-of-the-art methods, either because they contain excessive noise or because they violate one or more assumptions inherent to the chosen analysis method. To our knowledge, BNP-Step is the first time-series analysis algorithm which leverages Bayesian nonparametrics to learn the number and location of transitions between states and the emission levels associated to each state, while providing rigorous estimates of uncertainty for the learned quantities. We anticipate our algorithm will allow analysis of data sets at levels of noise or sparsity beyond what current state-of-the-art methods allow, and could potentially reveal previously unknown features in data sets analyzed using existing methods.

## INTRODUCTION

Time-series data appears everywhere from economics (1), to astronomy (2), to biophysics (3–5). Here our focus is on discrete state-space, discrete time transitions. That is, we consider systems hopping between different discrete states whose state is interrogated at discrete time intervals. We assume the states themselves are hidden. That is, when experiments report back on the state level, the state level is corrupted by noise.

For these data, two time-series data modeling paradigms exist: the Hidden Markov Models (HMMs) (6, 7) and step-finding methods, the latter of which are a subset of change-point detection methods (8, 9). Here we discuss the limitations of both in order to motivate a new tool.

HMMs are popular for analyzing noisy observations of stochastic sequences where a finite number of states are visited repeatedly (6, 7, 10). However, the standard HMM requires pre-specifying the number of states, which can be difficult for poorly-characterized systems (3, 10). The infinite HMM (iHMM) resolves this particular difficulty by using hierarchical Dirichlet nonparametric priors (11–13) to simultaneously estimate the number of states over the duration of the experiment, their transition points, and their emission characteristics (12, 14). However, the iHMM relies on states being revisited to reinforce learned parameters (12, 14), which can be problematic for data sets where some states are rarely or never revisited. For example, recent advances in single molecule force spectroscopy have revealed rare or hidden intermediate states in many biologically relevant molecules (15–20). Additionally, the expected number of states learned by an iHMM scales only logarithmically with the amount of data (14). This limitation can be circumvented by use of a Pitman-Yor process in place of a Dirichlet process (21, 22). However, both Dirichlet and Pitman-Yor processes pre-assume the form of the distribution of holding times: geometric in the case of the Dirichlet process, and discrete power law (such as the zeta or Zipf distributions) for the Pitman-Yor process (22). Examples of data sets with holding times that do not cleanly follow either of these distributions are found in a variety of fields, for example Biology (23), Chemistry (24), and Telecommunications (25).

An alternative class of methods—which do not suffer from the limitations of the HMM paradigm—termed step-finding methods rely on constrained optimization of a cost function. While many cost functions exist (9), here we focus on methods using information criteria (IC) (20, 26–29). These methods forgo a strict dynamical model in exchange for a different set of limitations (29, 30). In general, an information criterion consists of a log-likelihood term with a regularization to limit the number of state changes (*i*.*e*., steps) identified from the data. This regularization is drawn from information theoretic arguments in the case of the Akaike information criterion (AIC)(27), while for the Schwarz information criterion (also known as the Bayesian information criterion, or BIC) the regularization term originates from asymptotics on the posterior in the large data limit (26).

While these information criteria do not rely on dynamical models like HMMs to bias the identified location of state transitions, they still have myriad limitations: 1) models need to be compared head-to-head, which limits computationally efficiency (26, 28, 31); 2) they rely on greedy implementations (32–34) which can reduce accuracy (35, 36); and 3) they are deterministic and provide only point estimates (26, 27, 30).

In order to improve upon the limitations of both the IC and HMM paradigms, we propose BNP-Step, a Bayesian nonparametric (BNP) step-finding algorithm. From a time-series of scalar-valued observations, our method learns the number and time of transitions, the emission levels of the visited states, and their noise characteristics. In doing so, we seek to extend the scope of analyzable data beyond what current state-of-the-art step-finding permits in terms of noise and data sparsity, as shown in Figure 1. We demonstrate that BNP-Step outperforms both the iHMM (selected as the best representative of the HMM family) and BIC methods (selected as the best representative of the step-finding methods) on general data sets, and can successfully analyze data sets for which those algorithms fail. While there may be specific data sets for which our algorithm underperforms compared to the iHMM, these sets necessarily meet the assumptions of the iHMM where, for example, it is known that holding times are explicitly geometric and it is therefore preferred to bias the identification of state changes according to those expected statistics.

**Figure 1:**
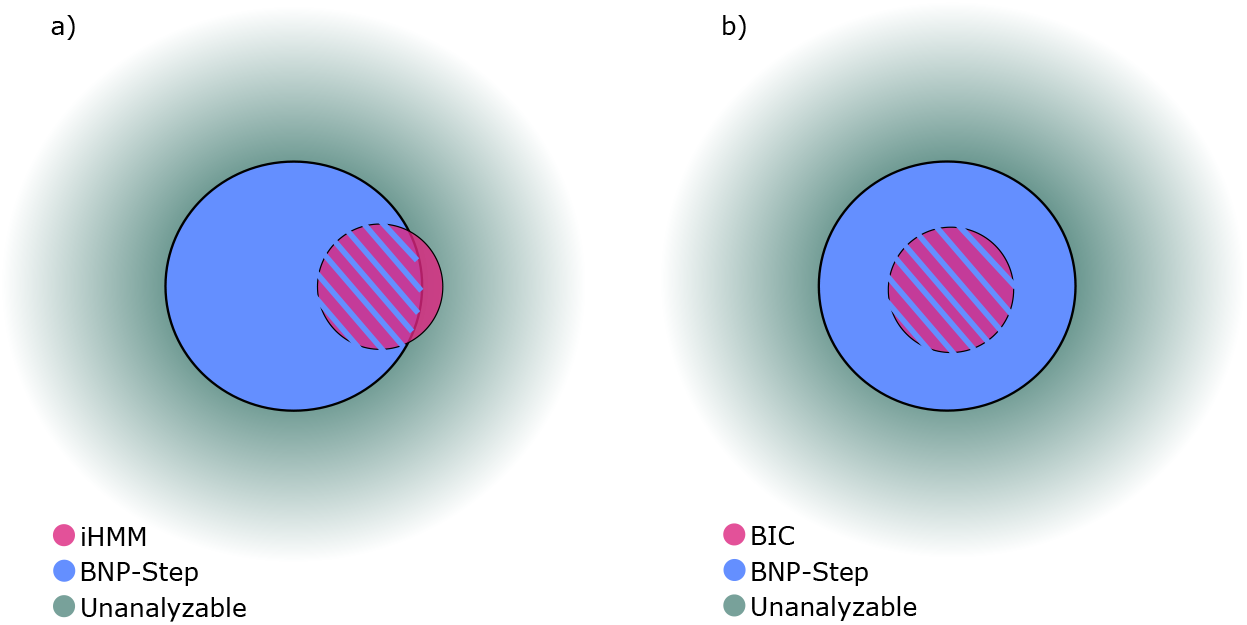
Illustration comparing the set of analyzable data sets for BNP-Step and two competing algorithms. (a) Illustration of the set of analyzable data sets for the iHMM and BNP-Step. The blue circle represents the set of data sets that BNP-Step can accurately analyze. The purple circle represents the set of data sets analyzable by an iHMM. The purple region with blue striations represents data sets that can be accurately analyzed by either method. The green space outside both circles represents data sets that are not currently analyzable by either method. (b) Illustration of the set of data sets analyzable by BNP-Step and BIC methods. The blue circle represents data sets analyzable by BNP-Step, while the purple region with blue striations represents the set of data sets analyzable by both BNP-Step and by BIC methods.

## METHODS

### Measurement model

In what follows, we define steps as discrete transitions occurring between two states with distinct emission levels. We assume data drawn as time-ordered scalar observations 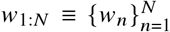 collected at times *t*_1:*N*_, where *N* is the total number of observations. We will connect these data to an observation model specifying the steps we wish to learn, the emission rates associated with each step, the time at which each step occurs, the background contribution, and the noise hyper-parameters, which we will collect into a parameter vector *θ*. Using Bayes’ theorem, we will then construct the posterior—a probability distribution over models conditioned on data—from which we will learn models alongside rigorous error estimates.

While the framework we propose is general, here, for sake of concreteness we focus on a Normal emission model. Under this assumption, as depicted in Figure 2, the observation associated to time *t*_*n*_ arises from the sum,

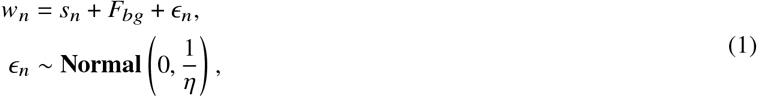

where *s*_*n*_ is the base emission level, *F*_*bg*_ is a constant offset bias term representing a background contribution constant with respect to time, and *∈*_*n*_ is the noise contribution assumed Normal. Here, we parameterize *∈*_*n*_ using a measurement precision *η*. Though we assume Normal noise here for ease of comparison with other methods, different noise models can be accommodated by changing the distribution giving rise to *∈*_*n*_ and adjusting the relevant prior and hyper-parameters to accommodate the new distribution.

**Figure 2:**
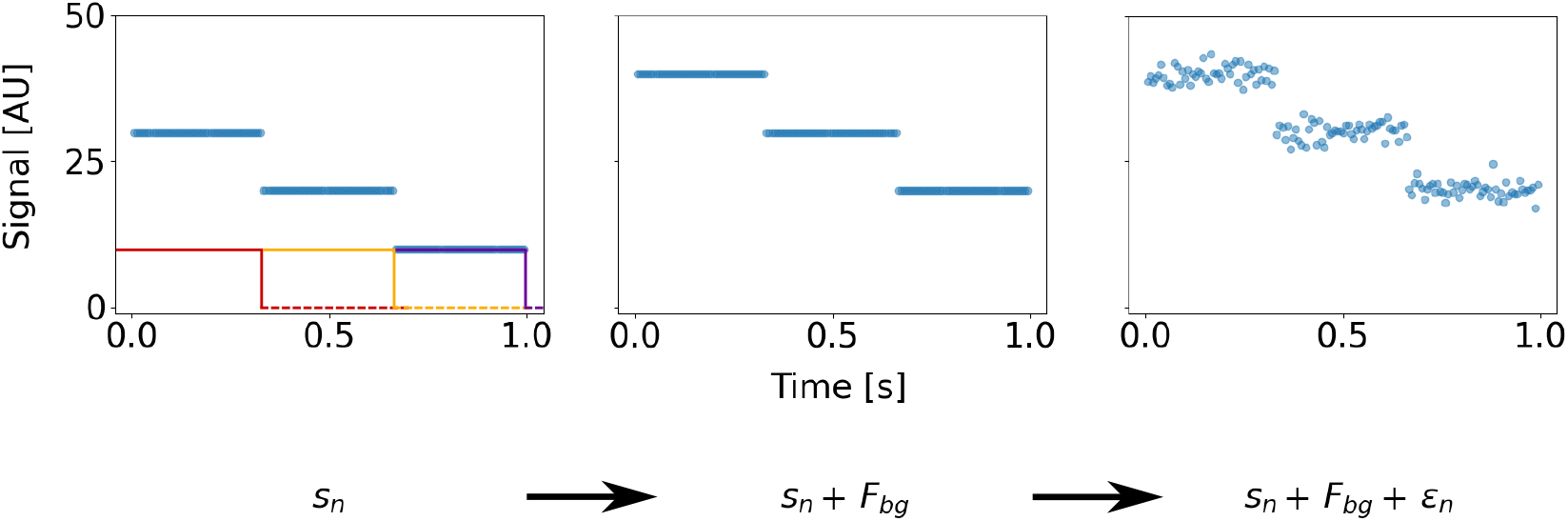
Illustration of contributions of *s*_*n*_, *F*_*bg*_, and *∈*_*n*_ **to observations** *w*_1:*N*_. For the purposes of demonstration only, we use a simple synthetic data set with three steps clearly visible by eye, where each interval between steps has a total of 50 associated observations, and where the constant background contribution is *F*_*bg*_ = 10.0 arbitrary units (AU). The left panel depicts the base emission level *s*_*n*_ associated with each observation, which is itself an infinite sum of scaled Heaviside functions. The first three of these terms are depicted by the red, yellow, and purple lines. All other terms in this sum (corresponding to steps associated to *b*_*m*_ = 0 terms) have been omitted as they contribute nothing. The middle panel depicts the addition of the constant background contribution *F*_*bg*_ to the base emission level of each observation. The right panel shows the addition of a Normal noise contribution with *η* = 0.5 to the sum of the base emission level and the background contribution, resulting in the observations *w*_1:*N*_ .

As the number of steps is unknown, we assume, within a Bayesian nonparametric paradigm that, *a priori*, the number of steps are infinite and ultimately decide on how many are warranted based on the data supplied. In other words, the base emission level *s*_*n*_ is itself a sum of contributions from an arbitrarily large number of possible steps, and can be expressed as

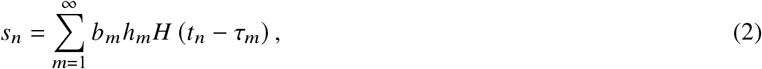

where *b*_*m*_ is a binary indicator variable comprising the nonparametric component of our model (outlined in the section titled “Model Inference” below and on which we place a Beta-Bernoulli process prior (37–39)), *h*_*m*_ is an emission rate that characterizes the *m*^*th*^ step, τ_*m*_ is the time at which the *m*^*th*^ step occurs, and *H* (*t*_*n*_ − τ_*m*_) is the step response function. We define *M* as the maximum possible number of steps in the system, which is theoretically infinite. A visualization of the contributions of each term to the base emission level at a given observation is shown in Figure 2. For compactness of notation, the step response function is taken to be a Heaviside function, which we detail in Section 1.1 of the Supplemental Information (SI).

With this observation model in hand, the likelihood reads

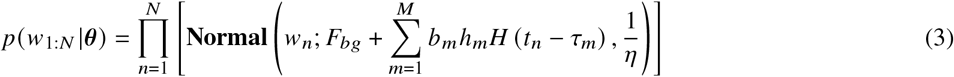

where *θ* =(*F*_*bg*_, *b*_1: *M*_, *h*_1: *M*_, τ_1: *M*_, *η*) collects all unknowns.

### Model inference

In order to determine the full posterior from the likelihood, we place appropriate priors on each of the unknown random variables gathered in *θ*. Since the number of steps—and therefore the number of required model parameters—is unknown *a priori*, we invoke the Bayesian nonparametric Beta-Bernoulli process prior (37, 38). That is, we place Bernoulli priors on each *b*_*m*_,

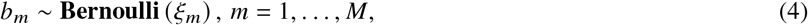

where the ξ_*m*_ are sampled from a Beta distribution. Following Ref. (40), we marginalize out the Beta prior, *i*.*e*., marginalize over ξ_*m*_, for which we find that the maximum number of active elements 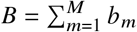 is binomially distributed (39, 41). To ensure convergence as *M* → ∞, we choose ξ_*m*_ = γ/ *M* (38, 41), where the new hyper-parameter γ tunes the mean number of assumed active *b*_*m*_ in the prior. Next, for computational ease, we make a finite approximation to the Beta-Bernoulli process prior by selecting a large albeit finite *M* (42). Typically, we select *M* ≤ *N* since two transitions that occur between neighboring observations result in an undetectable step. The remaining priors on quantities *F*_*bg*_, *h*_1: *M*_, τ_1: *M*_, and *η* are otherwise standard and are detailed in SI Section 1.2.

Since the full posterior does not attain an analytic form, we must draw samples from it using a custom Markov Chain Monte Carlo sampler wrapped in an overall Gibbs scheme (43, 44). Within our Gibbs scheme, when parameters, such as *F*_*bg*_, *b*_1: *M*_, *h*_1: *M*_, and *η* as we will see, admit analytic conditional posteriors we sample from them directly. Meanwhile, we sample parameters that do not have analytic conditional posteriors via a (brute-force) Metropolis algorithm (44, 45). A full description of all conditional posteriors and the methods used to sample them may be found in SI Sections 1.3 and 1.4.

### Data sources

To provide a standard basis for comparison between BNP-Step and other methods, we generated synthetic data sets according to the measurement model presented above. For comparison to the iHMM, two types of data sets were generated. In Type I data sets, the emission levels for each step were sampled from a bimodal Normal mixture distribution where the two means were separated by 3.5 standard deviations. Holding times were sampled from a multi-exponential distribution and discrete measurements were then generated by recording the emission level at regular time intervals. Each measurement was then corrupted with Normal noise. In Type II data sets, the system’s state space contained two frequently visited states and one rarely visited state. Data sets were generated and selected such that the rare state was visited only once in each trajectory. The holding times for these sets followed an exponential distribution, and each discrete measurement in these trajectories was generated in the same manner as described above for Type I sets. Our motivation for generating data sets with these characteristics was to simulate systems which seemingly satisfy the assumptions of the iHMM, but for which the iHMM gives misleading or incorrect results. In these cases, an iHMM may incorrectly attribute the subtle differences between states to noise.

For comparison to IC methods, of which the Kalafut and Visscher (KV) algorithm from Ref. (32) is a good example, and for general robustness testing, a third type of data set was generated (Type III). In these data sets, a fixed number of steps were generated, where the difference in base emission levels across each step—which we define as the step height—and the holding times were fixed at the same values throughout each data set. Individual observations were drawn from a Normal measurement model. A summary of all synthetic data set types and their relevant features is presented in Table 1.

**Table 1:** Data set types.

| Data Set Type | Emission Levels | Holding Times |
| --- | --- | --- |
| Type I | Normal-distributed | Multi-exponential |
| Type II | Fixed, 3 states | Exponential |
| Type III | Fixed, uniform step heights | Fixed duration |

To showcase BNP-Step’s performance on experimental data, we also analyzed data sets obtained from two optical tweezers experiments in Ref. (46). In the first experiment, the folding and unfolding of ss-DNA into single G-quadruplex (GQ) complexes was observed. In the second experiment, the elongation of telomere DNA by human telomerase was observed. Specific details on the characteristics of all data sets may be found in SI Sections 1.5 and 1.6.

## RESULTS AND DISCUSSION

### Comparison to iHMM

For comparison with BNP-Step, we used the implementation of the iHMM presented in Ref. (12). In Figure 3, we present representative trajectories from BNP-Step and from the iHMM for Type I data sets, at a range of signal-to-noise ratios (SNRs)—our SNR convention is given in SI Section 2. For BNP-Step, the learned trajectory corresponds to the maximum *a posteriori* (MAP) estimate, while for the iHMM it corresponds to the mode emission mean estimate trajectory, as defined in SI Section 2. While the learned trajectories for both methods predictably exhibit increasing deviation from ground truth as the SNR decreases, BNP-Step discriminates between small differences in emission levels at all SNRs (Figure 3), while the iHMM begins to attribute these differences to noise as the SNR decreases on account of states not being revisited.

**Figure 3:**
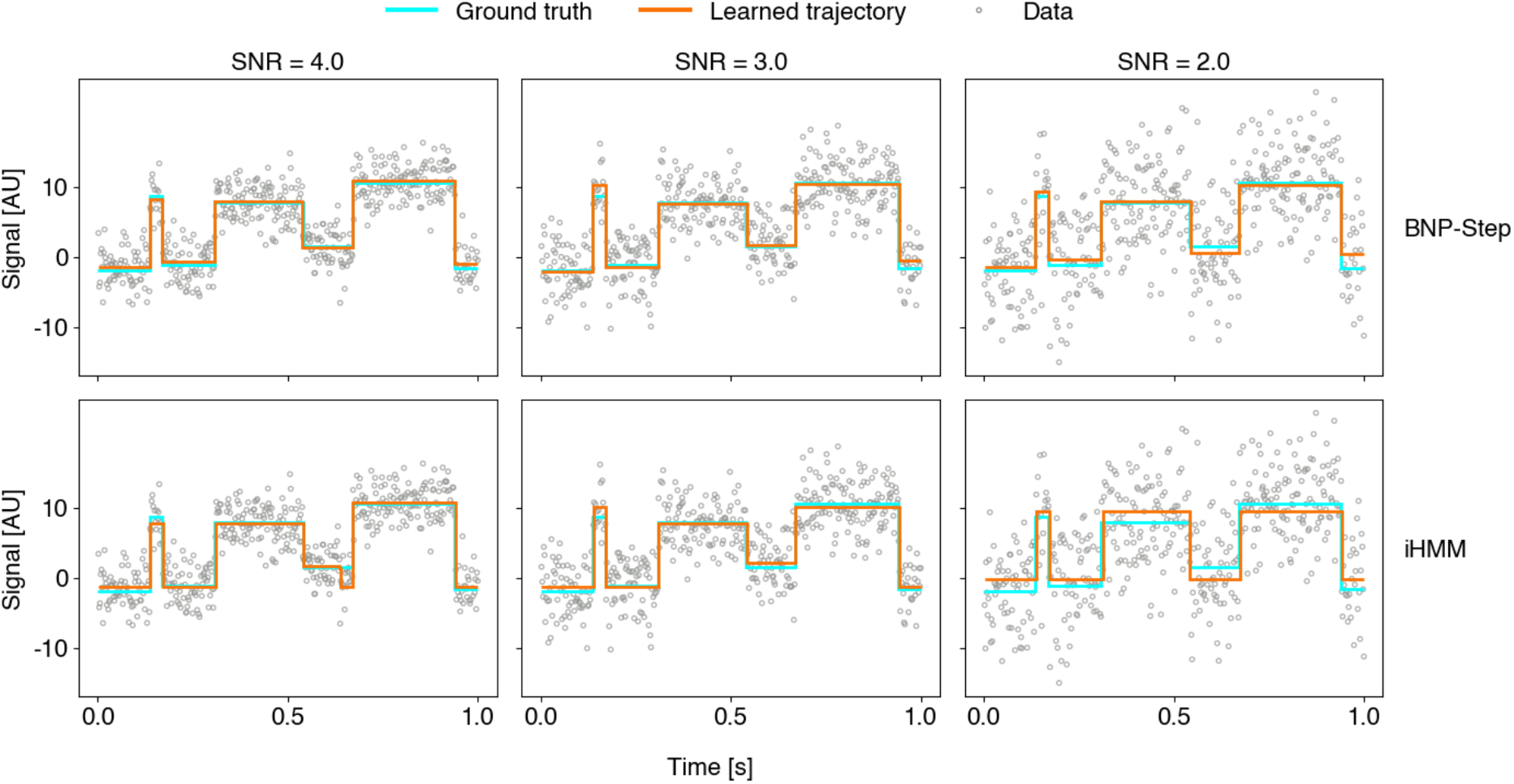
Trajectories learned by BNP-Step and the iHMM for a representative synthetic data set at SNR = 4.0, SNR = 3.0, and SNR = 2.0. The top row represents BNP-Step applied to a representative synthetic data set, while the bottom row represents the iHMM applied to the same data set for a given SNR. The cyan line, where visible, represents the ground truth trajectory, while the orange line represents each algorithm’s learned trajectory. The grey circles represent the raw synthetic data.

The histograms in Figure 4a show the joint posterior distributions of the state emission levels, conditioned on the mode number of states for the iHMM and the MAP number of transitions for BNP-Step, and marginalized over all other variables in *θ* —a detailed definition of these distributions is given in SI Section 2, and for brevity we will subsequently refer to these distributions as posteriors. As BNP-Step does not directly learn state emission levels, we reconstruct state emission levels for each sample by substituting the learned *b*_1: *M*_, *h*_1: *M*_, and τ_1: *M*_ into Equation 2, adding the learned *F*_*bg*_ to generate a piecewise constant trajectory of emission levels, then assigning the emission level of the holding period prior to the *m*^*th*^ step as the *m*^*th*^ state emission level. For the iHMM, even at SNR = 4.0 distinct peaks appear at some, but not all, of the ground truth values. What is more, as states do not repeat and therefore are never reinforced, the iHMM interprets the difference between states as noise and incorrectly merges them, resulting in the spurious peak between two states at SNR = 4.0. As the SNR decreases, the density around certain ground truth values approaches zero, and the peaks shift to regions between ground truth values, indicating inappropriate state merging. At the lowest SNR, the iHMM only learns two states, excluding *a posteriori* a significant portion of the ground truth signal levels. In contrast, since BNP-Step learns parameters associated with steps rather than states, it does not rely on the revisiting of states to reinforce its learned parameters and thus assigns posterior probability to all ground truth state emission levels, capturing their distributed nature. Figure 4b shows the posterior distributions of individual emission levels from BNP-Step, conditioned on the MAP number of transitions. At all SNRs, the ground truth values for all states lie within the 95% Credible Regions (CRs) generated from BNP-Step. As the SNR decreases, the posterior distributions for all states broaden, indicating increased uncertainty.

**Figure 4:**
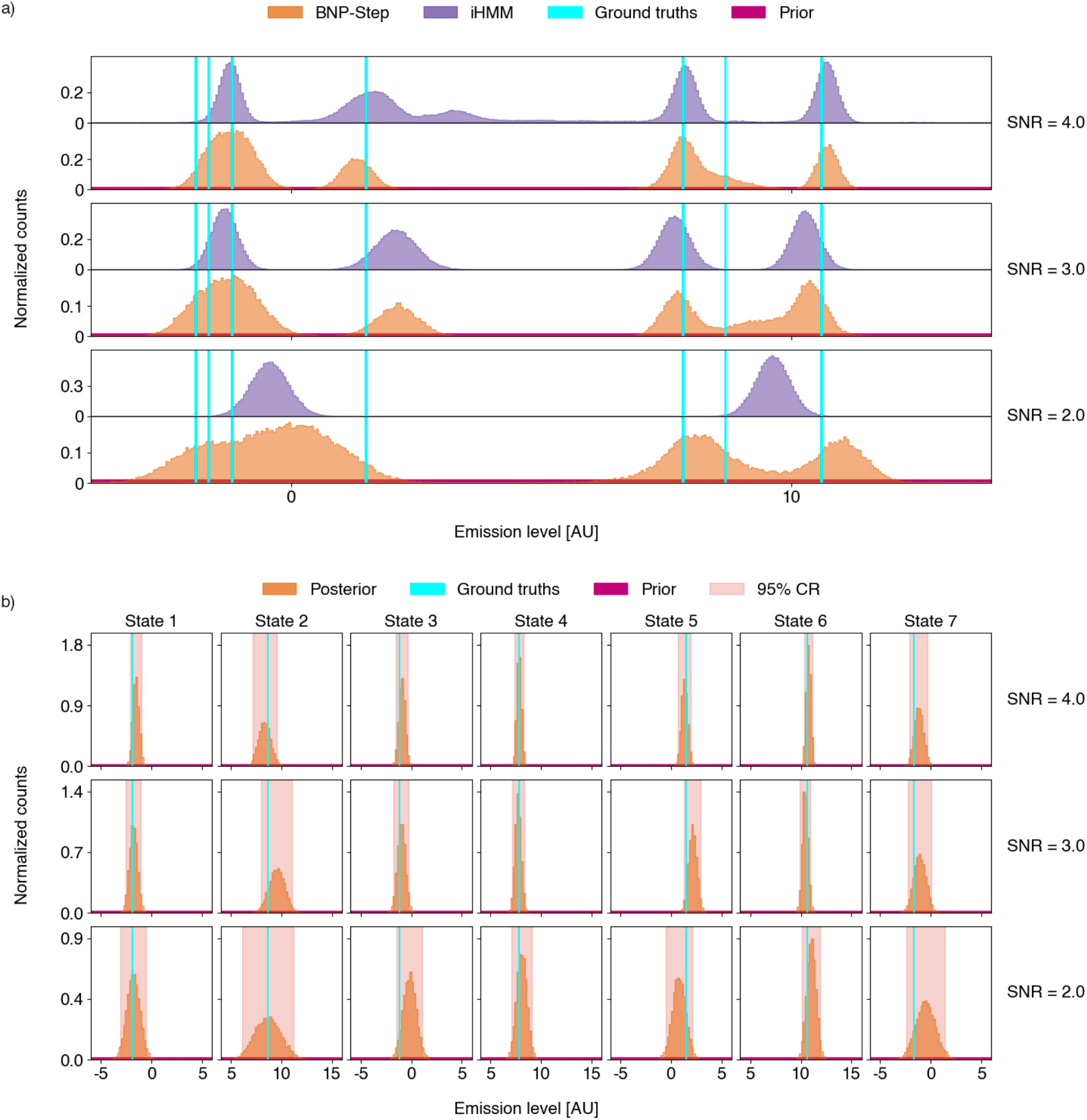
Conditional posterior distributions for SNR = 4.0, SNR = 3.0, and SNR = 2.0. a) Joint conditional posterior distributions for all state emission levels from BNP-Step and from the iHMM, as defined in SI Section 2. The samples from BNP-Step represented here were selected from the top 10^4^ MAP estimates after burn-in with the MAP number of steps for the data sets shown in Figure 3. The iHMM samples represent all samples after burn-in with the mode number of states. The cyan lines represent the common ground truth values for the emission levels of each state, while the magenta line represents the prior distribution utilized in BNP-Step. The height of BNP-Step’s posterior distributions compared to the prior indicates that our choice of prior has minimal effect on the learned values. b) Conditional posterior distributions from BNP-Step for the individual states in the data sets of Figure 3. The posteriors are marginalized over all other parameters in the same manner as described in SI Section 2. Each column represents one of the ground truth states in Figure 3, ordered chronologically. The pale red shaded areas represent the 95% Credible Regions (CRs).

Figure 5 shows the results of BNP-Step and the iHMM when applied to Type II data sets. Here, we expect the iHMM’s performance to improve relative to Type I data sets as the holding times are exponential, which satisfies one of the assumptions inherent to the HMM paradigm. However, the iHMM merges the two high-emission states into a single state on account of the rare state’s lack of reinforcement. This behavior is demonstrated qualitatively in the representative trajectories in Figure 5a. Notably, Figure 5b demonstrates that the iHMM is able to learn the emission levels of the two common states, further suggesting that lack of reinforcement of states is the reason behind the iHMM’s failures. Even *post hoc* tuning of the iHMM’s hyper-parameters to favor the recruitment of new states does not meaningfully improve its performance on these data sets, as shown in Figure S1 in SI Section 2.1. Meanwhile, BNP-Step produces three clear peaks centered around the ground truth values, indicating that it can successfully differentiate the rare state from the two common states.

**Figure 5:**
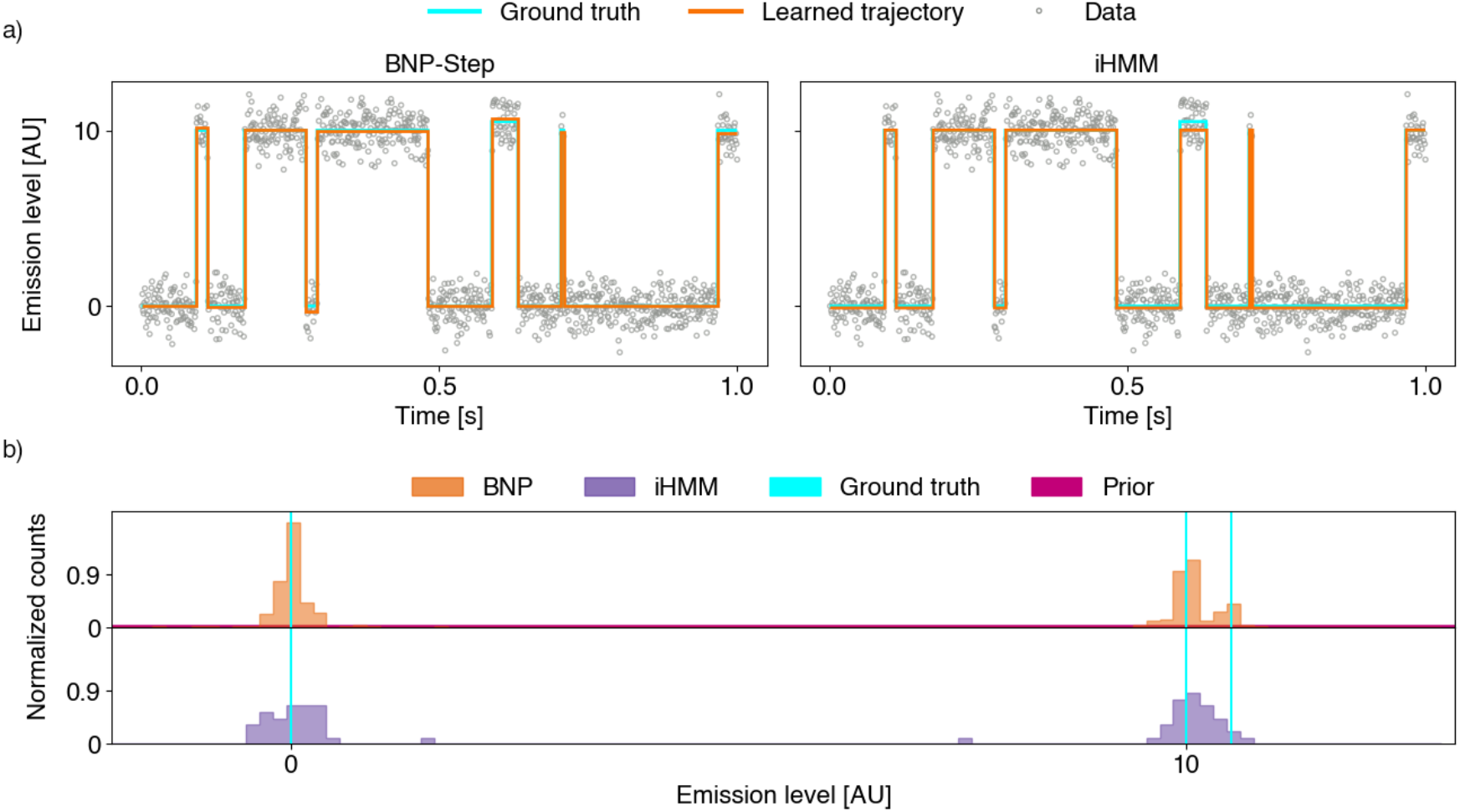
Analysis of three-state synthetic data by BNP-Step and the iHMM. a) Learned trajectories for BNP-Step and the iHMM, for a representative synthetic data set. The same labeling and color scheme is used as in Figure 3. b) Joint conditional posterior distributions (as described in SI Section 2) for all emission levels, conditioned on the mode number of samples (iHMM) or the MAP number of states (BNP-Step). For BNP-Step, the top MAP samples were selected from 30 synthetic data sets. For the iHMM, the mode emission levels as defined in SI Section 2 were used for each of the same 30 data sets. The same labeling and color scheme is used as in Figure 4.

For data sets with low noise, exponentially-distributed holding times, and repeatedly visited states, the iHMM outperforms BNP-Step, as evidenced in Figure S2 in SI Section 2.1. However, these data sets completely satisfy the inherent assumptions of the iHMM, while iHMMs are often applied for data where their assumptions hold only approximately (for example, when dynamics are assumed to be Markovian while transitions are actually due to multi-step biochemical reactions). We therefore conclude that when data clearly satisfies its assumptions, the iHMM is the obvious choice. However, for cases where it is not clear whether these assumptions are satisfied or they are satisfied only approximately, we argue that BNP-Step is less likely to return misleading or spurious results.

### Comparison to BIC-based algorithm

Figure 6 shows learned trajectories for both BNP-Step and the KV method for representative Type III data sets at a range of SNRs (with the SNR convention provided in SI Section 3). Again, both methods show generally decreasing agreement with ground truth as SNR decreases. However, as SNR decreases the KV method fails systemically, over-fitting noise as expected on account of its greedy nature, whereas BNP-Step merely becomes appropriately more uncertain at decreasing SNR, showing minimal over-fitting even for an SNR nearing unity.

**Figure 6:**
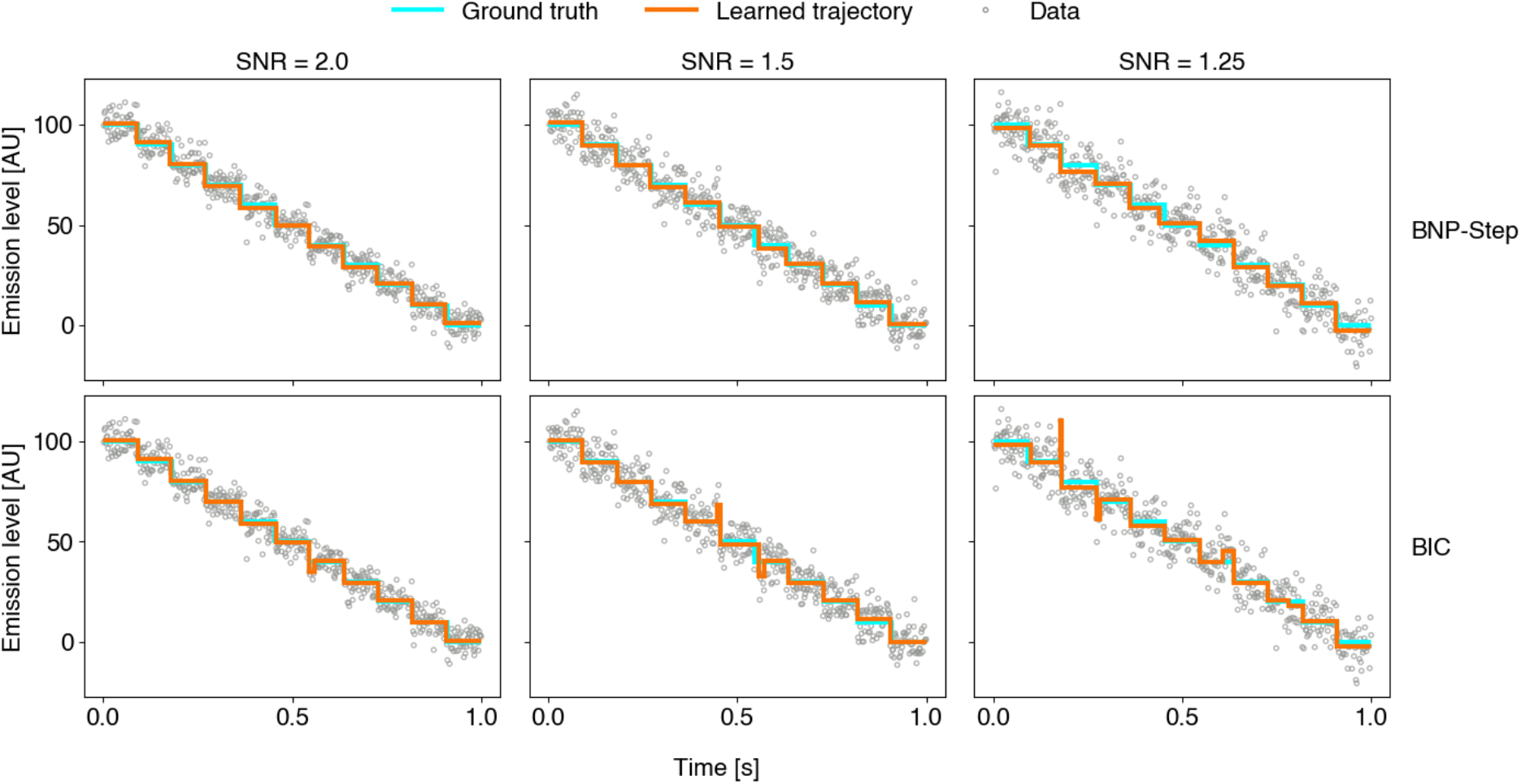
Learned trajectories for BNP-Step and the KV method, for representative synthetic data sets at SNR = 2.0, SNR = 1.5, and SNR = 1.25. The top row represents BNP-Step applied to a representative synthetic data set, while the bottom row represents the KV method applied to the same data set in each respective column. The same color scheme is used as in Figure 3. For BNP-Step, the trajectory corresponds to the MAP estimate, while for the KV method the trajectory corresponds to the point estimate returned by the algorithm.

Learned step height (as defined in the section titled “Data Sources”) and holding time histograms (Figures 7 and 8, respectively) demonstrate this behavior more clearly, where owing to the fixed nature of the step heights and holding times, we can generate frequentist histograms from the results of the KV method. As SNR decreases, both algorithms’ histograms become broader, indicating the previously noted loss of agreement with ground truth with increasing noise, but notably, at all SNRs tested, the KV method exhibits significantly broader distributions compared to BNP-Step for both step heights and holding times, indicating poorer performance in accurately learning these parameters. Additionally, the KV method’s holding time histograms begin to exhibit bimodal behavior at low SNRs, with the second peak centered around the inter-observation time of 30 ms. This indicates that KV begins to over-fit to individual data points at low SNRs. In contrast, BNP-Step consistently provides accurate unimodal distributions in which the ground truths are contained within the 95% CRs.

**Figure 7:**
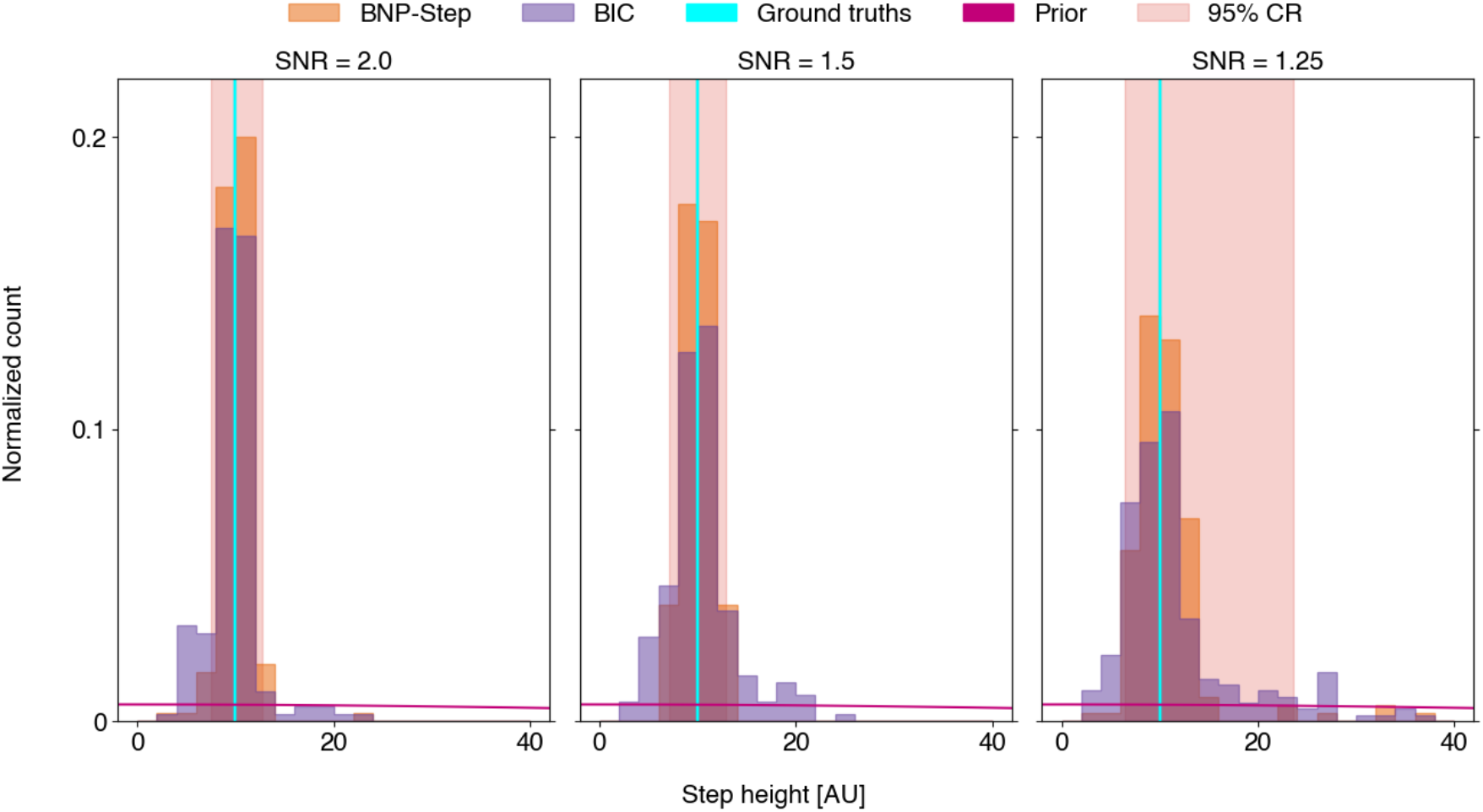
Normalized histograms of sampled step height magnitudes for both BNP-Step and the KV algorithm, for SNR = 2.0, SNR = 1.5, and SNR = 1.25. The orange bars represent the MAP estimates from BNP-Step for the analyzed synthetic data sets at each SNR, while the purple bars represent the KV algorithm’s point estimates for the same synthetic data sets. The cyan line represents the common ground truth value for the magnitude of the fixed step heights, while the magenta line represents the prior utilized in BNP-Step for the step heights. The sharp peaks of the histogrammed heights compared to the broad prior indicates that our choice of prior has a minimal effect on the learned value of these parameters. The pale red shaded region represents the 95% CR for BNP-Step.

**Figure 8:**
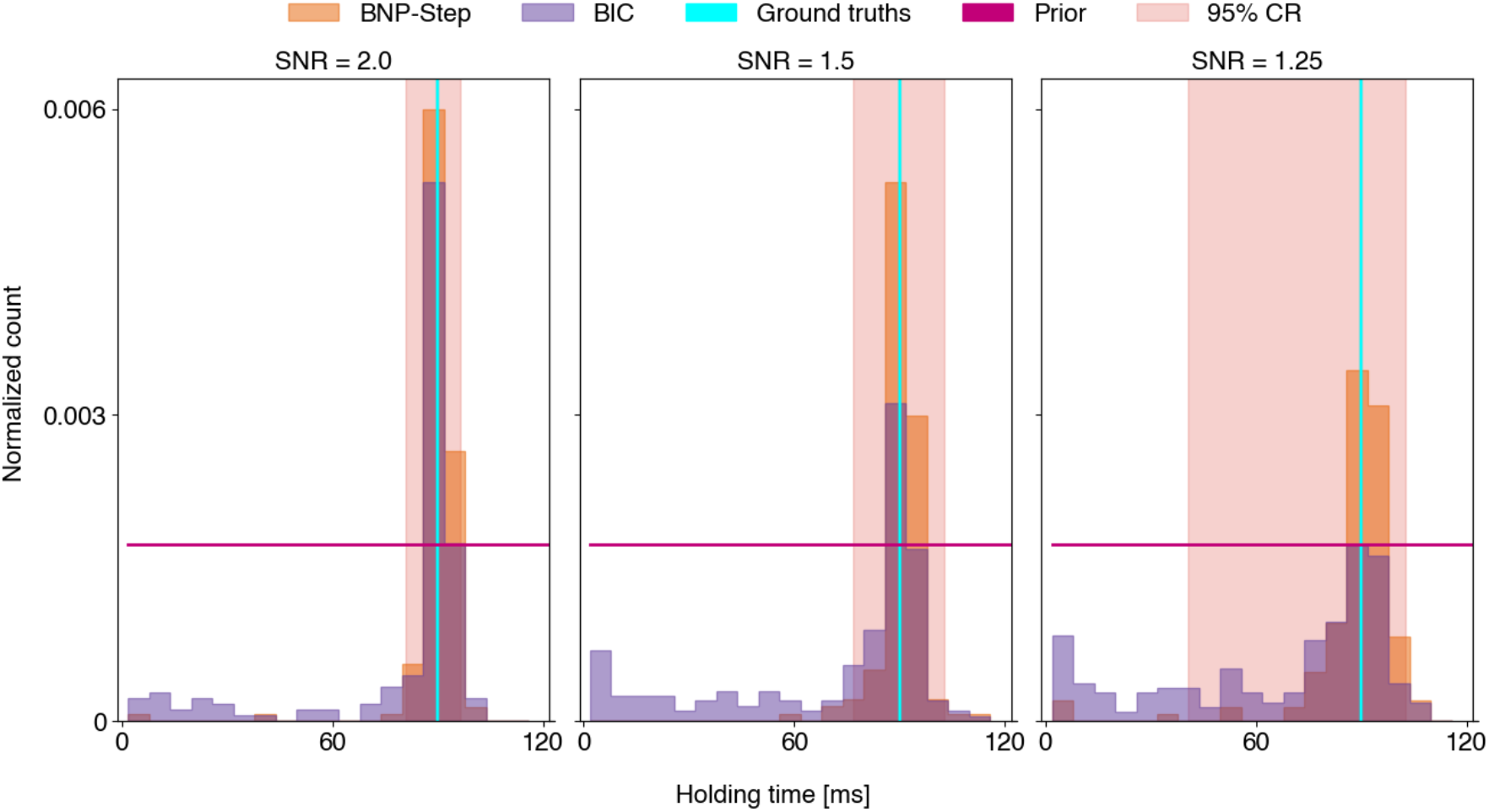
Normalized histograms of sampled holding times for BNP-Step and the KV algorithm, for SNR = 2.0, SNR = 1.5, and SNR = 1.25. The same labeling and color scheme is used as in Figure 7.

We also evaluated BNP-Step’s performance against the KV method as data density decreased (with data density defined in SI Section 3). These results mirror the results presented above almost exactly, so we have relegated the associated figures and discussion of them to SI Section 3.

### Application to experimental data

#### Single GQ-complex formation

Figure 9 shows representative trajectories for BNP-Step, the KV method, and the iHMM when applied to the ss-DNA GQ-complex data set described in the section titled “Data Sources”. Qualitatively, BNP-Step captures nearly all prominent features of the trajectory while ignoring small fluctuations at length scales below the physics of the experiment, whereas the KV method interprets some of these fluctuations as distinct steps. While BNP-Step misses three visually apparent short steps, two of these have data densities on par with the lower limit of BNP-Step’s characterized effectiveness (as described previously in the section titled “Comparison to BIC-based Algorithm” and in SI Section 3). Overall, the iHMM appears to perform comparably to BNP-Step in a qualitative sense, however we note that it required substantial manual tuning of both the transition and base concentration hyper-parameters to achieve this level of performance. Meanwhile, BNP-Step required only adjustment of the weak limit *M* to a larger value, reflecting the larger number of possible transitions in this much larger data set as compared to the synthetic data sets. While not strictly required for BNP-Step’s success, small upward adjustments of γ for large data sets (as shown in SI Table 1) resulted in improved time to convergence.

**Figure 9:**
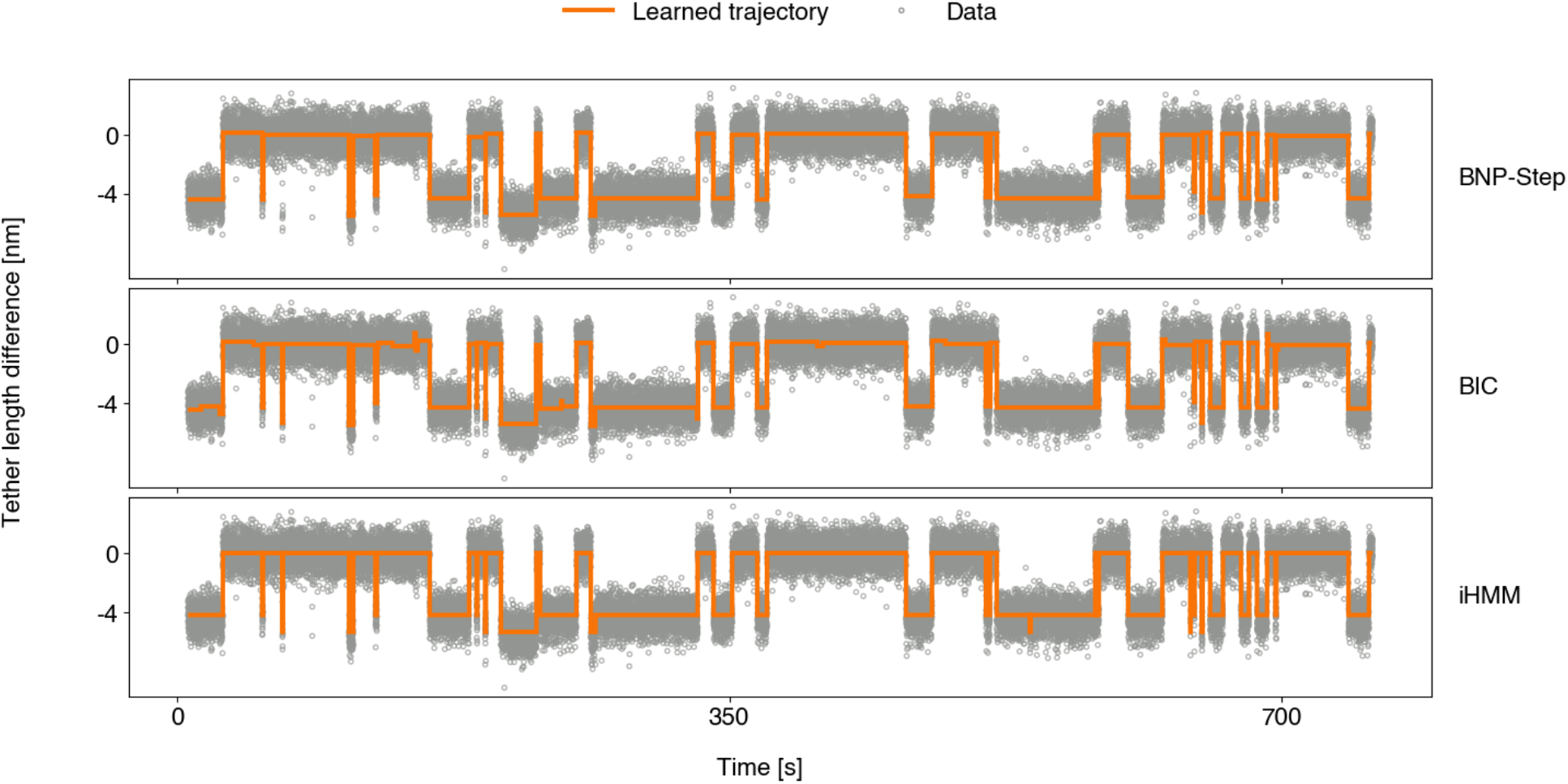
Learned trajectories for GQ-complex data set from BNP-Step, KV algorithm, and iHMM. The top panel represents BNP-Step applied to the GQ-complex data set, while the middle and bottom rows represents the KV method and the iHMM applied to the same data set, respectively. The same color scheme is used as in Figures 3 and 6. As in Figure 6, BNP-Step’s trajectory corresponds to the MAP estimate, while the KV method’s trajectory corresponds to its point estimate and the iHMM’s trajectory corresponds to the mode emission mean estimate. Here, tether length difference refers to the change in tether length with respect to the unfolded state.

Figure 10 shows the joint posterior distributions over all tether length changes from BNP-Step (conditioned on the MAP number of steps) and the iHMM (conditioned on the mode number of states) as described in SI Section 2, along with a frequentist histogram of the point estimates from the KV method. Both BNP-Step and the iHMM’s joint posterior distributions over the tether length changes show two distinct peaks near the length changes associated with two known GQ-complex extensions, denoted as ‘basket’ type and ‘hybrid-2’ type (46). While the peak associated with the larger of the two length changes is broader and not centered on the predicted value for both the iHMM and BNP-Step, this state was also substantially rarer. Notably, the iHMM also produces two clear peaks at smaller tether length differences, indicating that the iHMM assigns posterior probability to models that fit features at length scales substantially smaller than the known physics of the system. In contrast, while the KV method also produces two distinct peaks at the length changes predicted for GQ-folding, it also produces a large number of small steps consistent with over-fitting to noise on account of its greedy nature.

**Figure 10:**
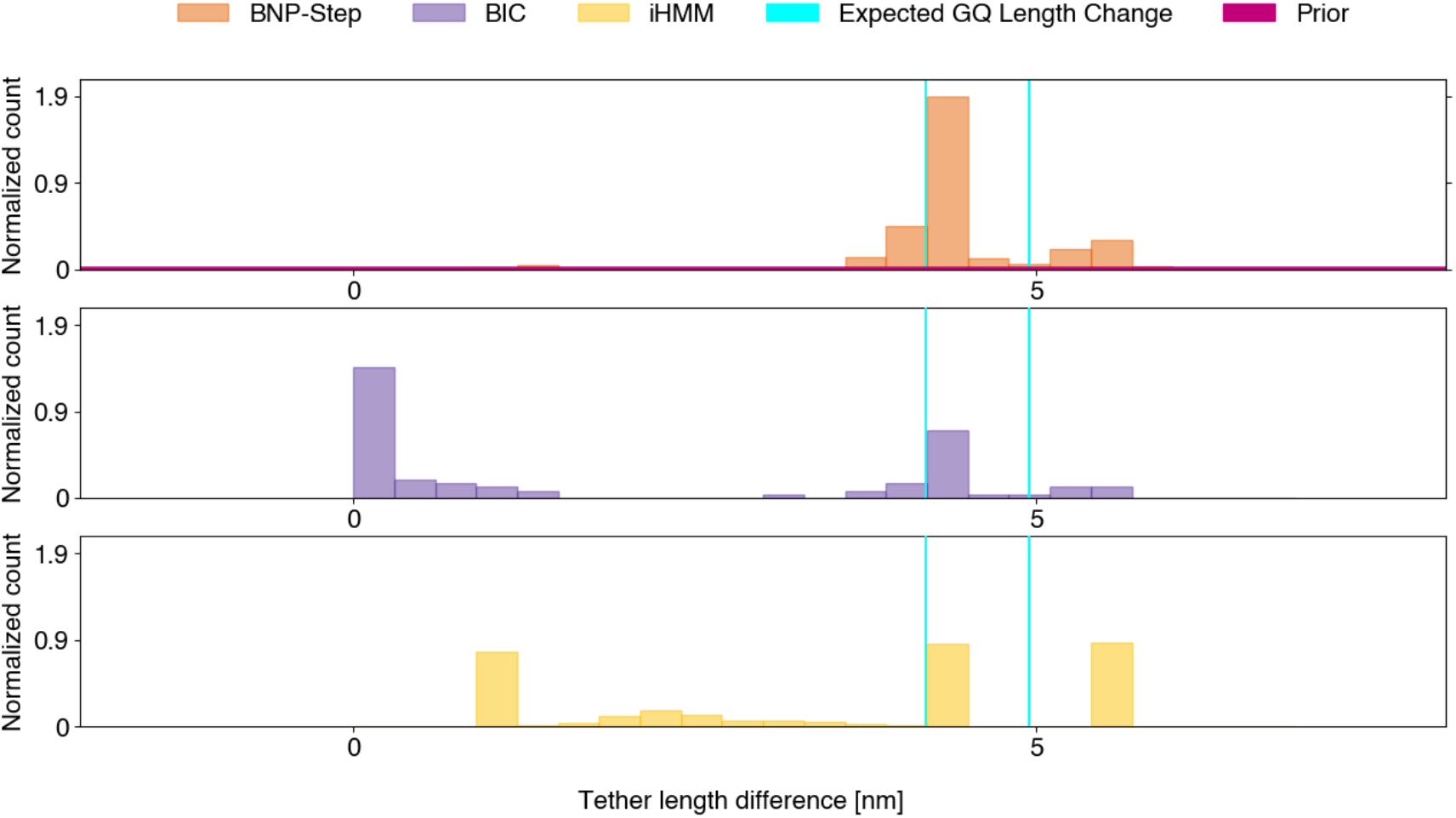
Joint posterior distributions of tether length changes for GQ-complex data from BNP-Step and the iHMM, with point estimate histogram of tether length changes learned by KV method. The same definition of the joint posterior as given in SI Section 2 is used. The orange bars represent all sampled tether length changes from the top 100 MAP estimates from BNP-Step (conditioned on the MAP number of steps), the purple bars represent the point estimate tether length changes learned by the KV algorithm, and the yellow bars represent all sampled tether length changes from the iHMM with the mode number of states. The cyan lines represent the predicted GQ tether length changes from Ref. (46). The magenta line represents the broad prior used in BNP-Step. As in Figure 9, tether length change refers to the change in tether length with respect to the unfolded state.

To compare our method’s results with those in Ref.(46), we present in Figure 11 survival curves (referred to as inverse cumulative distributions in Ref.(46)) for the holding times learned by BNP-Step. Figure 11a displays survival curves for holding times associated with GQ-unfolding events and GQ-folding events. Notably, the holding times for GQ-folding events appear to follow a single-exponential distribution, consistent with the findings in Ref.(46). In contrast, GQ-unfolding events do not exhibit a single-exponential distribution, as expected given the presence of two distinct folded emission levels (46). Furthermore, Figure 11b demonstrates that the holding times for basket type GQ-unfolding events—associated with the shorter of the two expected tether length changes—also deviate from a single-exponential distribution, suggesting the presence of multiple states associated with this emission level. These findings align with those in Ref. (46). Due to an insufficient number of hybrid-2 type GQ-complexes in the data set, a similar analysis of this emission level is omitted.

**Figure 11:**
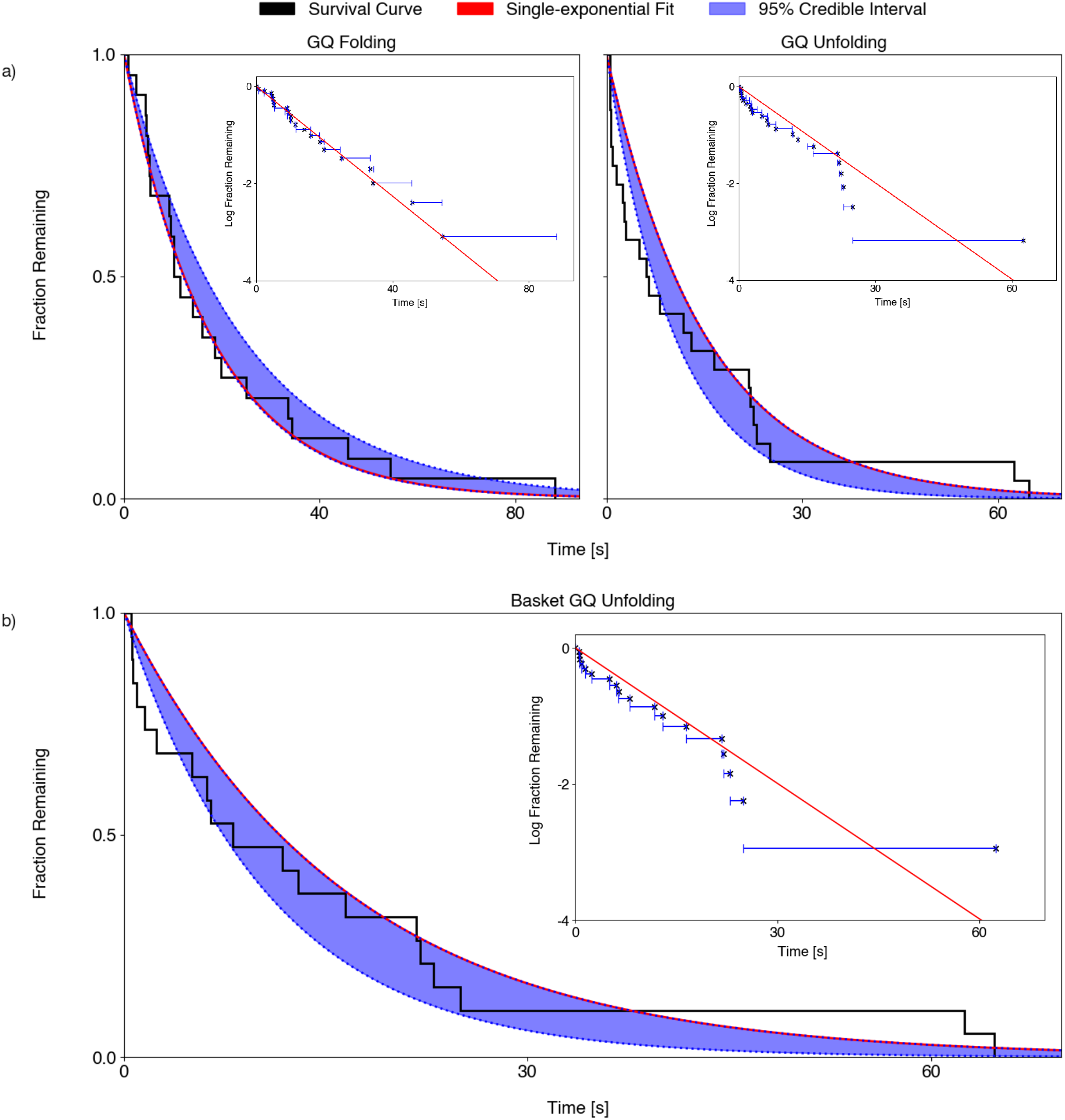
Survival curves for holding times for GQ-folding and unfolding events for the trajectory in Figure 9, with single-exponential fits. a) Survival curves for all GQ-folding events (left) and for all GQ-unfolding events (right). The black line represents the survival curve, while the red line represents the fit to a single exponential decay function generated using Numpy’s curve_fit function. Each inset shows the same survival curve and single-exponential fit in log space. b) Survival curve for basket GQ-unfolding events. The same labeling and color scheme is used as in part a.

#### Telomere DNA elongation

Figure 12 shows several representative trajectories learned by BNP-Step and the KV method for select telomere DNA elongation data sets from Ref. (46). Qualitatively, BNP-Step appears to capture the most prominent data features, although there are several areas where BNP-Step appears to be over-fitting continuous variations in the signal, as shown in inset (a) in Figure 12. However, the KV method also over-fits in these regions, and in addition learns a large number of very short steps which in some cases overwhelm visible features of the data, as shown in inset (b) in Figure 12. While Ref. (46) successfully analyzed these data sets with an iHMM that includes drift correction (13), we chose to compare BNP-Step to the iHMM used in previous sections—which does not include drift correction—to ensure a fair comparison as BNP-Step does not correct for drift. As this version of the iHMM produced visibly poor fits to these data sets, no further analysis was undertaken using this method. The iHMM’s learned trajectories may be found in Figure S6 in SI Section 4.

**Figure 12:**
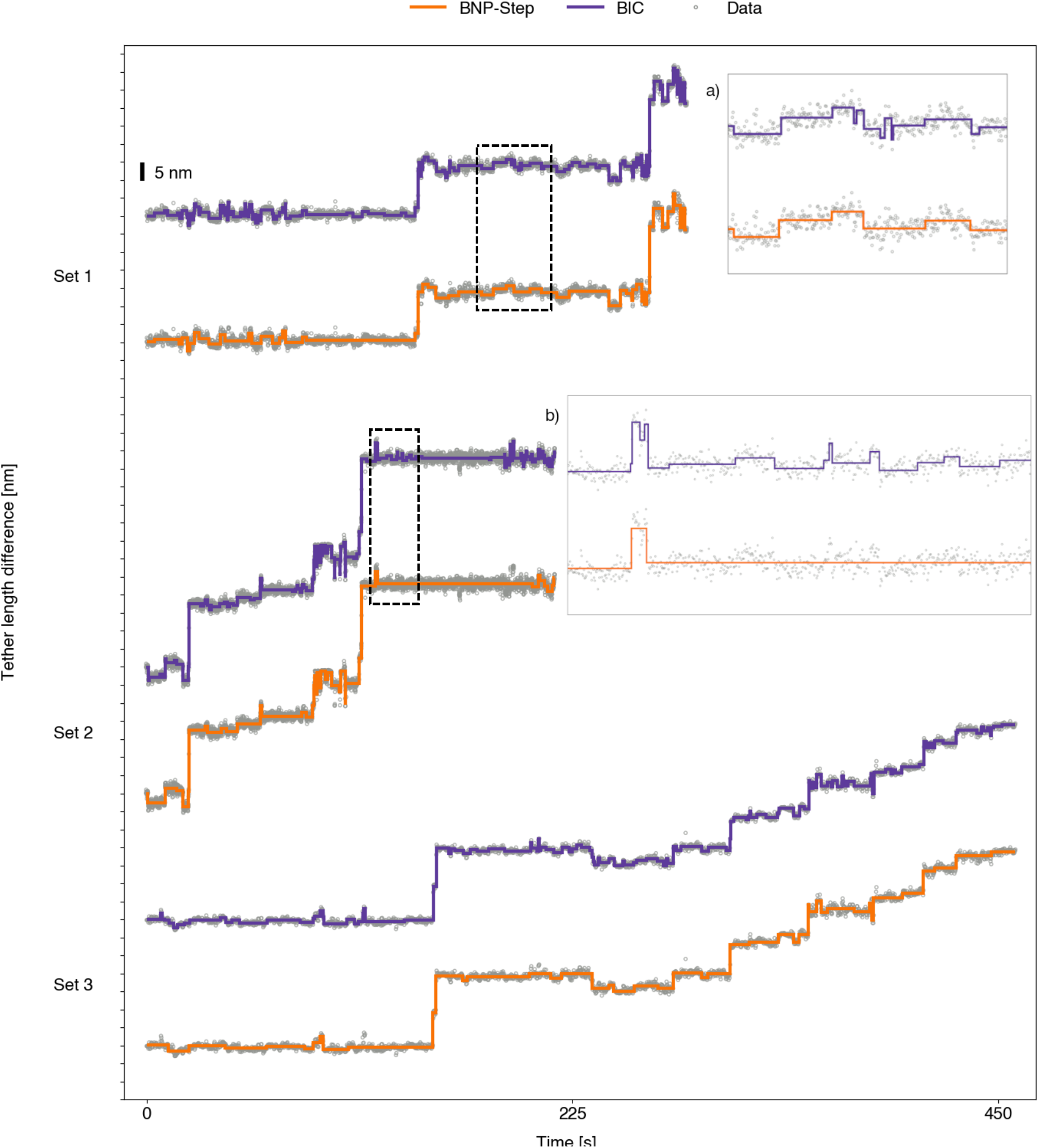
Representative fits to select telomerase DNA elongation data sets for BNP-Step and for the KV method. The orange line represents BNP-Step’s MAP estimate trajectory for each data set, while the purple line represents the point estimate trajectory for the KV method. Inset (a) shows a magnified view of a region (indicated by the dashed box) where both BNP-Step and the KV method appear to over-fit continuous signal features, while inset (b) shows a magnified view of a region where the KV method overfits severely while BNP-Step does not.

Figure 13 shows the joint posterior distributions (as defined in SI Section 2) of learned tether length changes from BNP-Step along with the distribution of the KV method’s point estimates of the tether length changes for data sets 1 and 3 as shown in Figure 12. In both cases, BNP-Step produces prominent peaks near 4.2 nm, which is the tether length change associated with formation of basket type GQ-complexes (46). While BNP-Step also produces peaks near 5.0 nm—the tether length change associated with hybrid-2 type GQ-complexes (46)—there appears to be a much greater degree of uncertainty associated with this feature, as the peak is shifted leftward in Set 1, and is very subtle and wide in Set 3. Histograms for data set 2 are shown in Figure S7 in SI Section 4. While there is a prominent peak near 4.2 nm in this data set as well, the peak nearest 5.0 nm is shifted substantially rightward, and thus is less likely to represent formation of hybrid-2 type GQ-complexes. Peaks associated with tether length changes greater than 5 nm are consistent with release of product DNA from an anchor site as described in Ref. (46). The irregular peaks located in the sub-4 nm region may represent over-fitting to low-frequency continuous variations in the signal, or uncharacterized motion of tether components. In contrast, the tether length changes learned by the KV method are overwhelmingly biased towards small steps below 2 nm, hindering its ability to learn larger steps.

**Figure 13:**
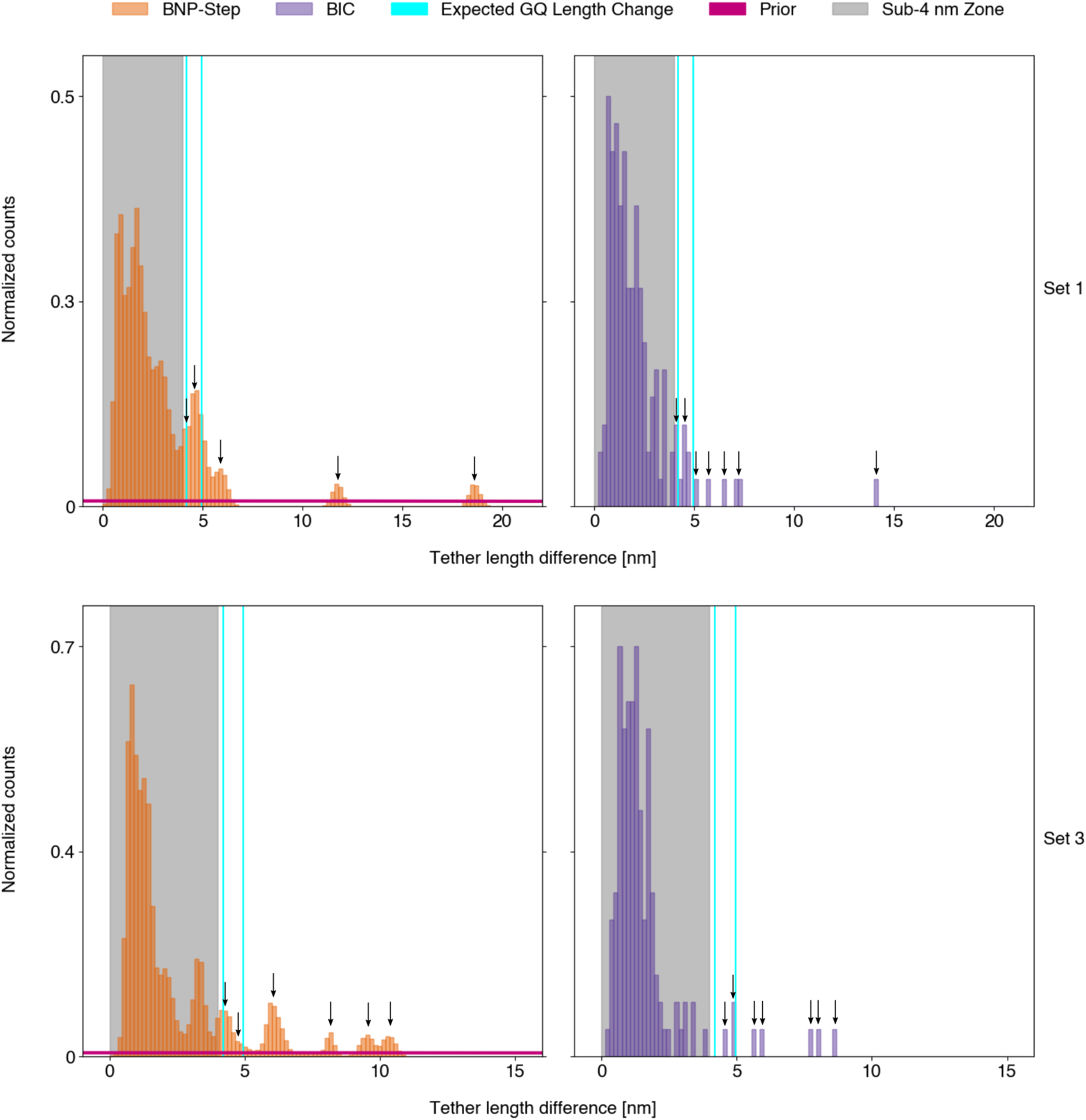
Joint posterior distribution of tether length changes from BNP-Step and point estimate frequentist histogram of tether length changes from the KV method, for telomere DNA elongation data sets 1 and 3 of Figure 12. We use the definition of the joint posterior as given in SI Section 2. The same color scheme is used as in Figures 7 and 8. For BNP-Step, the top 100 MAP samples (conditioned on the MAP number of steps) for each data set were used. Distinct peaks in the joint posterior distribution (BNP-Step) or frequentist histogram (KV method) correspond to distinct tether length changes learned by each method, and are indicated by the small black arrows. Peaks in the sub-4 nm region (shaded grey) are not marked by arrows owing to ambiguities in their origin.

### Robustness testing

To characterize BNP-Step’s robustness, we tested its performance at extreme values of the hyper-parameters *F*_*re f*_, *Ψ, h*_*re f*_, *χ, ϕ*, and γ, and at different ratios of step numbers to the weak limit *M* including 15 steps and *M* = 10, *M* = 30, and *M* = 50. In general, BNP-Step maintains acceptable performance for a broad range of hyper-parameters with the exception of the hyper-parameter γ in the Bernoulli priors on the *b*_*m*_, which should be set as low as feasibly possible since BNP-Step is biased towards fitting the maximum possible number of steps when γ approaches *M*. Our method also maintains performance for any *M* greater than or equal to the number of ground truth steps. Detailed information on all robustness tests may be found in SI Section 5, and Figures S7 and S8 in the SI quantify these results.

## CONCLUSION

BNP-Step represents a generalization of current state-of-the-art step finding techniques applicable to a broad variety of discrete space, discrete time data sets. In their formulation, iHMMs (indeed, virtually all HMM based methods) rely on states being re-visited in order to accurately learn their presence, their emission characteristics, and their connectivity to other states. However, if the states’ emission levels are distributed around some mean, or if there exist rarely visited states with characteristics similar to a more frequently visited state, then it may appear that states are revisited with emission levels that are uncertain *a posteriori*, when in reality states are not revisited at all. Essentially, small differences in the emission level are erroneously attributed to noise.

We demonstrated in the Comparison to iHMM section that for this reason, the iHMM fails to accurately learn the underlying trajectory and emission parameters for synthetic data sets with state emission levels varying around one of two mean values. As it does not depend on the revisiting of states for reinforcement, BNP-Step was able to successfully learn the emission parameters and trajectory while providing information about the uncertainty associated with each learned quantity. Even when presented with data sets that fulfilled one of the innate assumptions of the HMM paradigm—namely, exponential or geometric holding times—the iHMM still failed to identify the rarely-visited state in a three-state system, indicating that the revisiting of states is critical to the iHMM’s ability to learn trajectories and emission parameters. In contrast, BNP-Step was able to identify all three states.

Conversely, for data sets with a limited number of well-separated states visited repeatedly over the course of an experiment, BNP-Step learns slightly different emission parameters for all steps corresponding to a given state, a form of over-fitting to noise. However, we argue that such systems are obviously well-suited to analysis by iHMMs as they already conform to many of the assumptions inherent to iHMMs. For systems in which there exist a large, unknown number of states with similar emission characteristics, or in which there may be rarely-visited states with emission characteristics similar to more common states, BNP-Step is preferable as it is sensitive to subtle differences in emission parameters while also providing rigorous estimates of the uncertainty associated with all learned quantities.

BNP-Step can also analyze a wider variety of data sets than IC-based step-finding methods. For data sets with uniform step heights at low SNRs and low data densities, the KV method, a representative IC-based step-finding method, over-fits the data, placing spurious steps that often cannot be readily distinguished from real steps without *a priori* knowledge of the ground truth. This is a direct result of its greedy nature: once a step is detected by the method, it remains fixed, and the method is inherently unable to explore alternative models which lack this step. Furthermore, KV also provides only a point estimate of both the trajectory and the unknown parameters, which on their own are not enough to characterize the uncertainty over models. In contrast, BNP-Step’s use of the Beta-Bernoulli process allows it to explore the distribution of possible models by dynamically turning steps on and off in a probabilistic manner. This feature allows it to locate steps to a reasonable degree of accuracy at SNRs and data densities where IC-based methods fail, while simultaneously providing distributions for the heights and holding times for each step, allowing the calculation of error bars on the estimated parameters.

While we envision BNP-Step as being applicable to data sets across disciplines, we chose to demonstrate our algorithm’s effectiveness on biophysical data. When used on data sets from the optical tweezers experiments first presented in Ref. (46), BNP-Step confirms the presence of two distinct extensions of GQ-complexes. Additionally, BNP-Step is able to distinguish basket type GQ-complex formation in the telomere DNA elongation experiment despite the presence of low-frequency continuous variations in the data (commonly termed “drift”). In two of the three data sets analyzed, BNP-Step also offers possible evidence of hybrid-2 type GQ-complex formation, a subtle distinction not reported in the original paper (46), however as only a limited number of data sets were analyzed this finding is associated with a high degree of uncertainty. We also note that by failing to account for drift, we detect a potentially excessive number of small, transient steps in these data.

To our framework several additional generalizations can be made. As mentioned in the section titled “Measurement Model’, our method may be easily expanded to accommodate different noise models by replacing the additive, Normally-distributed *∈*_*n*_ and adjusting the associated priors. Finite instrument response times may be modeled by replacing the Heaviside function in the observation model with a sigmoidal function, whose parameters can be learned within the model. Different emission models (such as camera models for the case of biophysical data) may be accounted for by changing how the emission level is calculated.

For example, an integrative detector model can be accommodated by integrating the signal contribution over some pre-determined, finite exposure interval, potentially allowing for transitions occurring during an observation. Low-frequency drift may also be accounted for by allowing *F*_*bg*_ to vary continuously over the trajectory and sampling it using either a more expensive Gaussian Process prior or the cheaper but parametric spline parameterization (13). Additionally, while the current iteration of our algorithm assumes a constant precision throughout the experiment, precisions which change with each step may be accommodated by allowing *η* to have *M* components, each corresponding to a different holding period, and sampling each *η*_*m*_ separately.

For data sets with large numbers of steps, large numbers of observations, or both, computational cost must be taken into consideration. For a data set with *N* observations, BNP-Step scales as *0* (*M*^2^*N*) . Practically, for a data set of approximately 3.6 × 10^3^ observations where *M* = 100, BNP-Step generated 10^4^ samples in a wall time of approximately 1 hour on a machine with an Intel Core i7-11700 processor (16 x 2.5 GHz) and 16 GB DDR4 RAM. The performance of our algorithm may be further improved by use of a collapsed Gibbs sampling scheme (47), where we first marginalize over either *η* or *F*_*bg*_ and *h*_1: *M*_, then sample parameters *b*_1: *M*_ or τ_1: *M*_, then recover *η* or *F*_*bg*_ and *h*_1: *M*_ through direct sampling. While the computational cost of each step would increase, overall convergence would be faster as fewer samples would be required for convergence (47).

Our method extends IC methods’ reach without the confining model-based assumptions inherent to the HMM paradigm. To date, we are not aware of any other method leveraging BNPs to resolve differences in state emission levels small compared to the noise distribution while accurately detecting the time at which steps occur.

## DATA/CODE AVAILABILITY

Raw synthetic data, experimental data sets, and Python code for this study will available via **Github** upon editorial acceptance and review.

## AUTHOR CONTRIBUTIONS

S.P. and I.S. conceived and designed the method. A.R. derived key expressions and implemented the method. M.J.C. provided all experimental data and offered insights into its interpretation. A.R. produced synthetic data and analyzed all data sets. A.R., M.S., and S.P. wrote the article. A.R., M.S., M.J.C., I.S., and S.P. edited the article.

## ACKNOWLEDGMENTS

We thank W. Xu, A. Saurabh, J. S. Bryan IV, M. Fazel, and P. Pessoa for interesting discussions and insights. M.J.C. acknowledges support from NSF (MCB-1919439). S.P. acknowledges support from NIH NIGMS (R01GM130745), NIH MIRA (R35GM148237) and NIH NIGMS (R01GM134426).

## SUPPORTING CITATIONS

(48–52) appear in the supporting material.

## SUPPLEMENTARY MATERIAL

An online supplement to this article can be found by visiting BJ Online at http://www.biophysj.org.

