## Supplemental Information for "An accurate probabilistic step finder for time-series analysis"

#### 1 MATHEMATICAL DERIVATIONS

##### 1.1 Step response function

For ease of computation, we model an instantaneous step response using a Heaviside function,

$$H(t) = \begin{cases} 1, & t \le 0 \\ 0, & t > 0. \end{cases} \tag{S1}$$

Given times $t_{1:N}$ associated with observations $w_{1:N}$, and step times $\tau_{1:M}$, we can define a set of functions of the form $H(t_n - \tau_m)$. For a given observation occurring at $t_n$, emission rates $h_m$ associated with steps ending at times $\tau_m$ later than or equal to $t_n$ fully contribute to the base emission level, while those ending at times earlier than $t_n$ contribute nothing.

##### 1.2 Priors

On each unknown random variable, we place the following priors:

$$F_{bg} \sim \textbf{Normal}\left(F_{ref}, \frac{1}{\psi}\right), \tag{S2}$$

$$b_m \sim \textbf{Bernoulli}\left(\frac{\gamma}{M}\right); \; m = 1, \ldots, M, \tag{S3}$$

$$h_m \sim \textbf{Normal}\left(h_{ref}, \frac{1}{\chi}\right); \; m = 1, \ldots, M, \tag{S4}$$

$$\tau_m \sim \textbf{Uniform}_{[\tau_1, \tau_N]}; \; m = 1, \ldots, M, \tag{S5}$$

$$\eta \sim \textbf{Gamma}\left(\phi, \eta_{ref}\right), \tag{S6}$$

where the Uniform prior of Equation S6 is discretized over the times $t_{1:N}$. The prior on each $b_m$ originates from a Beta-Bernoulli process (1–3), in which the hyper-parameter associated with the Beta prior has been marginalized out for computational reasons as detailed in Ref. (4). The hyper-parameters $F_{ref}$, $\psi$, $h_{ref}$, $\chi$, $\phi$, and $\eta_{ref}$ were chosen to yield broad prior distributions. As discussed in Supplemental Section 5, we choose a low value for $\gamma$ to simulate observations taken on a time scale much shorter than the dynamics of the system. To avoid uncontrollable overestimation of the number of steps as $M$ tends to infinity, we emphasize that $\gamma$ must be kept independent of $M$. Note that $\gamma$ need not be assigned an integer value, although we chose integers to maintain the physical interpretation that $\gamma$ indicates the *a priori* expected number of steps. Numerical values for all hyper-parameters can be found in Table S1.

##### 1.3 Derivation of the full posterior and conditional posteriors

Armed with the likelihood,

$$p(w_{1:N}|F_{bg}, b_{1:M}, h_{1:M}, \tau_{1:M}, \eta) = \prod_{n=1}^{N}\left[\textbf{Normal}\left(w_n; F_{bg} + \sum_{m=1}^{M} b_m h_m H(t_n - \tau_m), \frac{1}{\eta}\right)\right], \tag{S7}$$

and the priors (Equations S2 through S6), we can construct an expression that is proportional to the full posterior,

$$p\left(F_{bg}, b_{1:M}, h_{1:M}, \tau_{1:M}, \eta | w_{1:N}\right) \propto \left[\prod_{n=1}^{N} p\left(w_n | F_{bg}, b_{1:M}, h_{1:M}, \tau_{1:M}, \eta\right)\right] p\left(F_{bg}\right) \left[\prod_{m=1}^{M} p\left(b_m\right) p\left(h_m\right) p\left(\tau_m\right)\right] p(\eta). \tag{S8}$$

Table S1: Numerical values for all hyper-parameters used for BNP-Step.

| Hyper-parameter | Value |
|---|---|
| $F_{ref}$ | 0.0 |
| $\psi$ | 0.00028 |
| $h_{ref}$ | 0.0 |
| $\chi$ | 0.00028 |
| $\phi$ | 1.0 |
| $\eta_{ref}$ | 10.0 |
| $\gamma_{M=20}$ | 1 |
| $\gamma_{M=50}$ | 2 |
| $\gamma_{M=100}$ | 5 |
| $\gamma_{M=200}$ | 5 |

In the limit $M \longrightarrow \infty$, because of the priors on $b_{1:M}$, the posterior converges (3–6). Hence, for sufficiently large $M$, the posterior conclusions are independent of $M$. For this reason, we specify an $M \gg 1$ in advance and never change it again. Our goal is to draw samples from the posterior using a Gibbs algorithm, for which we need the conditional posteriors for each unknown random variable. We describe the derivation of each conditional posterior in subsequent sections.

##### 1.3.1 Conditional posterior on $F_{bg}$, $h_{1:M}$

We will show that in generating a conditional posterior on both $F_{bg}$ and $h_{1:M}$ simultaneously, we can utilize conditional conjugacy to obtain an analytic form. Utilizing Bayes' theorem, we obtain

$$p\left(F_{bg}, h_{1:M} | w_{1:N}, b_{1:M}, \tau_{1:M}, \eta\right) \propto \left[\prod_{n=1}^{N} p\left(w_n | F_{bg}, b_{1:M}, h_{1:M}, \tau_{1:M}, \eta\right)\right] p\left(F_{bg}\right) \left[\prod_{m=1}^{M} p\left(h_m\right)\right]. \tag{S9}$$

Writing out the right-hand side using the appropriate probability density functions (PDFs) for the likelihood and the priors, we obtain

$$\begin{aligned} &p\left(F_{bg}, h_{1:M} | w_{1:N}, b_{1:M}, \tau_{1:M}, \eta\right) \propto \\ &\left(\frac{\eta}{2\pi}\right)^{N/2} \exp\left[-\frac{\eta}{2}\sum_{n=1}^{N}\left(w_n - F_{bg} - \sum_{m=1}^{M} b_m h_m H\left(t_n - \tau_m\right)\right)^2\right] \times \\ &\left(\frac{\chi}{2\pi}\right)^{M/2} \exp\left[-\frac{\chi}{2}\sum_{m=1}^{M}\left(h_m - h_{ref}\right)^2\right] \left(\frac{\psi}{2\pi}\right)^{1/2} \exp\left[-\frac{\psi}{2}\left(F_{bg} - F_{ref}\right)^2\right]. \end{aligned} \tag{S10}$$

Because they do not depend on $F_{bg}$ or any of the $h_{1:M}$ and because the right-hand side is proportional to the left-hand side, the terms in front of each exponential may be absorbed into an implied normalization constant. The remaining exponential terms may then be combined into a single exponential, which we will subsequently demonstrate has the form of an $(M+1)$-dimensional multivariate Normal PDF.

After combination of all exponential terms, the exponent—which we label here as $S$—has the form

$$S = -\frac{\eta}{2}\sum_{n=1}^{N}\left(w_n - F_{bg} - \sum_{m=1}^{M} b_m h_m H\left(t_n - \tau_m\right)\right)^2 - \frac{\chi}{2}\sum_{m=1}^{M}\left(h_m - h_{ref}\right)^2 - \frac{\psi}{2}\left(F_{bg} - F_{ref}\right)^2. \tag{S11}$$

Expanding the squared terms then yields

$$
\begin{aligned}
S = -\frac{1}{2}\Bigg[ \eta \Bigg( \sum_{n=1}^{N} \Bigg( & w_n^2 - 2F_{bg} w_n - 2w_n \sum_{m=1}^{M} b_m h_m H\left(t_n - \tau_m\right) + \left(F_{bg}\right)^2 \\
& + 2F_{bg} \sum_{m=1}^{M} b_m h_m H\left(t_n - \tau_m\right) + \left( \sum_{m=1}^{M} b_m h_m H\left(t_n - \tau_m\right) \right)^2 \Bigg)\Bigg) \\
& + \chi \left( \sum_{m=1}^{M} \left( \left(h_m\right)^2 - 2h_m h_{ref} + \left(h_{ref}\right)^2 \right) \right) \\
& + \psi \left( \left(F_{bg}\right)^2 - 2F_{ref} F_{bg} + \left(F_{ref}\right)^2 \right) \Bigg].
\end{aligned}
\tag{S12}
$$

All terms in the exponent that carry no dependence on $F_{bg}$ or any of the $h_{1:M}$ result in an exponential factor that can be safely absorbed into the normalization constant, leaving terms that are either linear or quadratic in $F_{bg}$ or one of the $h_{1:M}$, or are cross-terms.

A general multivariate Normal distribution has an exponent of the form

$$
-\frac{1}{2}\left(\overline{x} - \overline{\mu}\right)^T \Lambda \left(\overline{x} - \overline{\mu}\right), \tag{S13}
$$

where $\overline{x}$ is a vector of observations, $\overline{\mu}$ is the mean vector, and $\Lambda$ is the precision matrix. This may be expanded into a double sum,

$$
-\frac{1}{2} \sum_{i=1}^{M+1} \left[ \left(\overline{x} - \overline{\mu}\right)_{1i}^T \left[ \sum_{j=1}^{M+1} \Lambda_{ij} \left(\overline{x} - \overline{\mu}\right)_{j1} \right] \right] = -\frac{1}{2} \sum_{i=1}^{M+1} \sum_{j=1}^{M+1} \left[ x_{i1}\Lambda_{ij}x_{ji} - x_{i1}\Lambda_{ij}\mu_{ji} - \mu_{i1}\Lambda_{ij}x_{ji} + \mu_{i1}\Lambda_{ij}\mu_{ji} \right]. \tag{S14}
$$

As the fourth term in the brackets lacks dependence on any of the $x_{ij}$, we may safely ignore it as its counterpart in $S$ was absorbed into the normalization constant. Comparing this to $S$, we can readily extract the components of the precision matrix by matching the terms quadratic in the $h_{1:M}$ and $F_{bg}$ as well as the cross-terms. We then obtain the components of the precision matrix,

$$
\Lambda_{11} = \eta N + \psi, \tag{S15}
$$

$$
\Lambda_{i1} = \Lambda_{1i} = \eta \sum_{n=1}^{N} \left( b_i H\left(t_n - \tau_i\right) \right), \; i \neq 1, \tag{S16}
$$

$$
\Lambda_{ii} = \eta \sum_{n=1}^{N} \left( b_i H\left(t_n - \tau_i\right) \right)^2 + \chi, \; i \neq 1, \tag{S17}
$$

$$
\Lambda_{ij} = \eta \sum_{n=1}^{N} \left( b_i b_j H\left(t_n - \tau_i\right) H\left(t_n - \tau_j\right) \right), \; i \neq j, \; i \neq 1, \; j \neq 1. \tag{S18}
$$

Similarly, we can extract the components of $\overline{\mu}$ by matching terms linear in the $h_{1:M}$ and $F_{bg}$ between Equations S12 and S14. In doing so, we generate a set of linear equations,

$$
\begin{pmatrix}
\Lambda_{1,1} & \Lambda_{1,2} & \dots & \Lambda_{1,j} & \dots & \Lambda_{1,M+1} \\
\Lambda_{2,1} & \Lambda_{2,2} & \dots & \Lambda_{2,j} & \dots & \Lambda_{2,M+1} \\
\vdots & \vdots & & \vdots & & \vdots \\
\Lambda_{M+1,1} & \Lambda_{M+1,2} & \dots & \Lambda_{M+1,j} & \dots & \Lambda_{M+1,M+1}
\end{pmatrix}
\begin{pmatrix} \mu_1 \\ \mu_2 \\ \vdots \\ \mu_{M+1} \end{pmatrix}
=
\begin{pmatrix} q_1 \\ q_2 \\ \vdots \\ q_{M+1} \end{pmatrix}, \tag{S19}
$$

where the components $q_i$ are defined as

$$
q_1 = \eta \sum_{n=1}^{N} w_n + \psi F_{ref}, \tag{S20}
$$

$$q_i = \eta \sum_{n=1}^{N} \left( w_n b_i H\left(t_n - \tau_i\right) \right) + \chi h_{ref},\ i = 2 \ldots M + 1. \tag{S21}$$

We may therefore calculate $\overline{\mu}$ by inverting the precision matrix, then right-multiplying $\overline{q}$ with it.

While straightforward, this construction for the conditional posterior on $F_{bg}$ and $h_{1:M}$ comes with a high computational cost. If $M$ is large, this results in a large matrix that must be inverted in order to generate samples for $F_{bg}$ and $h_{1:M}$. However, from Equations S15 through S18 above we see that if $b_m = 0$, the corresponding element of the precision matrix is zero if $i \neq j$ and $\chi$ if $i = j$. Consequently, we may reduce the computational cost of sampling $F_{bg}$ and $h_{1:M}$ by inverting only the portion of the precision matrix corresponding to terms for which $b_m = 1$.

Using elementary row and column swaps, it is possible to rearrange the precision matrix such that all terms for which $b_m = 1$ are in a square block in the upper left quadrant, while those for which $b_m = 0$ and $i = j$ are along the main diagonal in the lower right quadrant; all other entries would be zero. We may then block-wise invert this matrix. However, since the lower left and upper right blocks are zero matrices, this amounts to finding the inverses of only the upper left and lower right blocks. Because the lower right block is diagonal, with all diagonal entries equal to $\chi$, its inverse is straightforward and may be deduced without needing to explicitly compute it. Consequently, we need only perform the more expensive inversion operation on the upper left block matrix corresponding to terms where $b_m = 1$.

Furthermore, since for $b_m = 0$ the $q_i$ reduce to $\chi h_{ref}$, the mean vector reduces to $h_{ref}$ for these terms. Therefore, we are justified in sampling the $h_m$ for which the associated $b_m$ is zero directly from the prior.

##### 1.3.2 Conditional posteriors on $b_m$

We may write the conditional posteriors on each $b_{1:M}$ as

$$p\left(b_m | \eta, F_{bg}, b_{-m}, h_{1:M}, \tau_{1:M}, w_{1:N}\right) \propto \left[ \prod_{n=1}^{N} p\left(w_{1:N} | \eta, F_{bg}, b_{1:M}, h_{1:M}, \tau_{1:M}\right) \right] p\left(b_m\right), \tag{S22}$$

where $b_{-m}$ represents the variables not currently being sampled. In terms of the PDFs, this is

$$p(b_m | \eta, F_{bg}, b_{-m}, h_{1:M}, \tau_{1:M}, w_{1:N}) \propto$$
$$\exp\left[ -\frac{\eta}{2} \sum_{n=1}^{N} \left( w_n - \left( F_{bg} + \sum_{m'=1}^{m-1} b_{m'} h_{m'} H(t_n - \tau_{m'}) + b_m h_m H(t_n - \tau_m) + \sum_{m''=m+1}^{M} b_{m''} h_{m''} H(t_n - \tau_{m''}) \right) \right)^2 \right] \left( \frac{\gamma}{M} \right). \tag{S23}$$

Since $b_m$ is a binary variable, this conditional posterior amounts to a Bernoulli distribution. We define the conditional posterior probability that $b_m = 1$ as Equation S23 with $b_m = 1$, normalized by the sum of this term with the value of Equation S23 when $b_m = 0$.

However, as the number of observations $N$ grows large, numerical underflow becomes a concern. To address this, we perform sampling in log-space. Since the normalization factor introduces a sum of exponentials, we may use the Gumbel trick (7) to facilitate computation of this term.

##### 1.3.3 Conditional posteriors on $\tau_m$

The conditional posteriors on $\tau_{1:M}$ may be written as

$$p(\tau_m | \eta, F_{bg}, b_{1:M}, h_{1:M}, \tau_{-m}, w_{1:N}) \propto \left[ \prod_{n=1}^{N} p(w_{1:N} | \eta, F_{bg}, b_{1:M}, h_{1:M}, \tau_{1:M}) \right] p(\tau_m), \tag{S24}$$

where $\tau_{-m}$ represents the $\tau_m$ not currently being sampled. Since our goal is to sample a specific $\tau_m$, all terms which do not depend on the $\tau_m$ in question may be absorbed into the implied normalization constant. All remaining terms in the exponent will depend on some power of $b_m h_m H\left(t_n - \tau_m\right)$. If the associated $b_m$ is zero, the exponent will vanish and no terms depending on $\tau_m$ will remain. Thus, for the case where the associated $b_m = 0$, the conditional posterior is a re-scaling of the prior. As a result, when $b_m = 0$ we sample $\tau_m$ directly from the prior.

When $b_m = 1$, no such simplification occurs and the conditional posterior on $\tau_m$ does not have an analytic form. In these cases we sample the conditional posterior using the Metropolis algorithm with proposals drawn from the prior (8).

##### 1.3.4 Conditional posterior on $\eta$

The conditional posterior on $\eta$ is given by

$$p(\eta|F_{bg}, b_{1:M}, h_{1:M}, \tau_{1:M}, w_{1:N}) \propto \left[\prod_{n=1}^{N} p(w_n|F_{bg}, b_{1:M}, h_{1:M}, \tau_{1:M}, \eta)\right] p(\eta). \tag{S25}$$

Written in terms of the PDFs, this corresponds to

$$p\left(\eta|F_{bg}, b_{1:M}, h_{1:M}, \tau_{1:M}, w_{1:N}\right) \propto$$
$$\left(\frac{\eta}{2\pi}\right)^{N/2} \exp\left[-\frac{\eta}{2}\sum_{n=1}^{N}\left(w_n - F_{bg} - \sum_{m=1}^{M} b_m h_m H\left(t_n - \tau_m\right)\right)^2 - \frac{\eta\phi}{\eta_{ref}}\right]\left(\frac{\phi}{\eta_{ref}\Gamma\left(\phi\right)}\right)\left(\frac{\eta\phi}{\eta_{ref}}\right)^{\phi-1}. \tag{S26}$$

As the Normal distribution is conditionally conjugate to the Gamma distribution, the conditional posterior for $\eta$ is a Gamma distribution with updated hyper-parameters $\alpha$ and $\beta$, where

$$\alpha = \frac{N}{2} + \phi, \tag{S27}$$

$$\beta = \left(\frac{\phi}{\eta_{ref}} + \frac{1}{2}\sum_{n=1}^{N}\left(w_n - F_{bg} - \sum_{m=1}^{M} b_m h_m H\left(t_n - \tau_m\right)\right)^2\right)^{-1}. \tag{S28}$$

#### 1.4 Sampling

The initial samples for $F_{bg}$, $h_{1:M}$, $\tau_{1:M}$, and $\eta$ were drawn from their priors. For the $b_{1:M}$, one of two strategies were used based on the characteristics of the data sets being analyzed. For the Type I and Type III synthetic data sets described in Supplemental Sections 1.5.1 and 1.5.2, respectively, the initial $b_{1:M}$ were sampled directly from the Bernoulli priors. For the Type II data sets described in Supplemental Section 1.5.1 and for all experimental data sets, all of the $b_{1:M}$ were initialized to 1 as this was empirically noted to help speed convergence. For all synthetic data sets, $2 \times 10^5$ samples were generated. For experimental sets, samples were drawn until either a total of $2 \times 10^5$ samples was reached, or the algorithm had run for 7 days, whichever event happened first.

To reduce burn-in time, we implemented a simulated annealing strategy (9, 10). The initial temperature and temperature schedule were chosen such that the temperature reached 1.0 at approximately $2 \times 10^4$ samples. All samples prior to burn-in were discarded prior to analysis.

#### 1.5 Synthetic data set generation

##### 1.5.1 Comparison to iHMM

Synthetic data sets were generated by directly implementing the observation model specified in Section 2.1 of the main text. For both types of data sets (as described in Section 2.3 of the main text), the base emission levels were constrained to alternate between one or more 'high' values and a 'low' value of 0.0 AU (arbitrary units), with the background emission level set to zero. For Type I data sets, the base high value was set to 10.0 AU. A Normally-distributed factor with a mean of zero and a variance of 2.0 was then added to the base value in order to generate state emission levels that varied around their respective base values. The holding times of each state followed a multi-exponential distribution with rate parameters $\lambda_1 = 3.3 \times 10^{-5}$ ms$^{-1}$, $\lambda_2 = 6.7 \times 10^{-5}$ ms$^{-1}$, and $\lambda_3 = 1.0 \times 10^{-4}$ ms$^{-1}$. Ten such base data sets were generated with 500 observations each, where the discrete observations were generated assuming a sampling frequency of one sample every 30 ms. Each observation was then individually corrupted with Normal noise to generate data sets with signal-to-noise ratios (SNRs) of 4.0, 3.0, and 2.0, where the SNR is defined in Supplemental Section 2.

For Type II data sets, instead of sampling the state emission levels from Normal distributions, a rare third state was added with an emission level of 10.5 AU. The holding times for all transitions for these sets were Exponentially-distributed with transition matrix

$$\Pi = \begin{pmatrix} 0.99 & 0.009 & 0.001 \\ 0.01 & 0.99 & 0.0 \\ 0.01 & 0.0 & 0.99 \end{pmatrix}, \tag{S29}$$

where the first row corresponds to the 0.0 AU state, the second row to the 10.0 AU state, and the third row to the 10.5 AU state, while the first, second, and third columns represent transitions to the 0.0 AU, 10.0 AU, and 10.5 AU states, respectively. Candidate data sets were selected by hand such that the rare state was visited only once in the trajectory. For the purposes of demonstrating the iHMM's success mode, one set where the rare state was never visited was selected. For all of the data sets described above, a weak limit of $M = 100$ was used. Additionally, for the purposes of figure generation, all data sets were subsequently re-scaled in time such that the total time for each trajectory was 1 second.

##### 1.5.2 Comparison to KV Method

For comparison to the method described in Ref. (11), the same general implementation of the observation model was used as in Supplemental Section 1.5.1. Each synthetic data set (described as Type III in Section 2.3 of the main text) contained 10 total steps and 11 total states, with emission levels decreasing by 10.0 AU after each step and a fixed 1500 ms holding time for each state. $M$ was set to 20, and the background emission level was set to zero. The variance of the Normal noise term was chosen such that the data set had the desired SNR as described in Supplemental Section 3. The time scales of these sets were also re-scaled for figure generation in the same manner as described in Supplemental Section 1.5.1.

##### 1.5.3 Robustness testing

For the purposes of robustness testing, synthetic data sets were generated in the same manner as those in Supplemental Section 1.5.2. A 5-step data set with a holding time of 360 ms for each state was generated to examine the behavior of the algorithm at varying values of the hyper-parameter $\gamma$. A 15-step set with a holding time of 1500 ms per step was generated for the purposes of testing varying values of the weak limit $M$. All sets were re-scaled in the same manner as described in Supplemental Section 1.5.1 for the purposes of figure generation.

#### 1.6 Experimental data sets

In order to evaluate our algorithm's performance on real experimental data, we sourced data sets from two optical tweezers experiments presented in Ref. (12). The first experiment followed the folding and unfolding of single-stranded DNA into a single G-quadruplex complex. Prior to analysis, the provided data set was de-drifted as specified in Ref. (12). The second experiment measured the elongation of telomere DNA by human telomerase. Candidate data sets were trimmed to exclude times prior to placement of the experimental apparatus into the reaction buffer, as well as tether breakage when this feature was present.

### 2 COMPARISON TO IHMM: SUPPLEMENTAL SECTION

When comparing BNP-Step to the iHMM presented in Ref (13), we define the SNR differently depending on the type of synthetic data set being analyzed (as described in Supplemental Section 1.5.1 above). For Type I data sets, we define the SNR as the ratio of the difference between the two base emission levels to the standard deviation of the Normal noise. For Type II data sets, we define the SNR as the ratio of the difference between the emission levels of the two common states (0.0 AU and 10.0 AU) to the standard deviation of the Normal noise. For both types of data sets, the iHMM's transition concentration hyper-parameter $\alpha$ and its base concentration hyper-parameter $\gamma$ were set to 1.0.

For results produced by the iHMM, trajectories were selected using the MATLAB function chainer_analyze_means from the SI of Ref (13), which generates the mode emission mean estimate trajectory of the time series. That is, it produces the most likely sequence of mean values of the emission distributions which produced the observations.

To compare the iHMM's and BNP-Step's estimates of the state emission levels, we generated joint conditional posterior distributions over all state emission levels, with all other parameters in $\boldsymbol{\theta}$ marginalized out. For the iHMM, we conditioned on the mode number of states—that is, the most likely number of distinct states—while for BNP-Step we conditioned on the MAP number of transitions. Additionally, as BNP-Step does not directly estimate state emission levels, we reconstructed them from the estimated parameters as described in the main text.

#### 2.1 Supplementary Figures

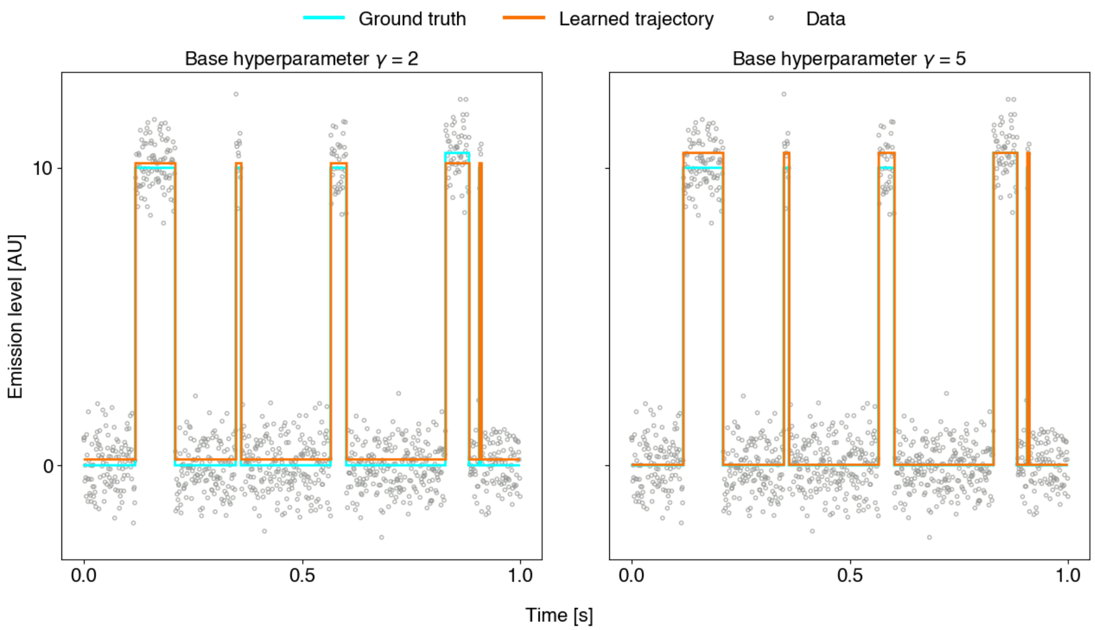


Figure S1: **Representative learned trajectories from the iHMM for three-state synthetic data, at base concentration hyper-parameter values $\gamma$ = 2 and $\gamma$ = 5 and SNR = 5.0.** The cyan line, where visible, represents the ground truth trajectory, while the orange line represents the mode emission mean estimate as described in Supplemental Section 2. The grey circles represent the raw synthetic data. Note the lack of improvement in performance even as the base concentration hyper-parameter is adjusted to favor the recruitment of additional states.

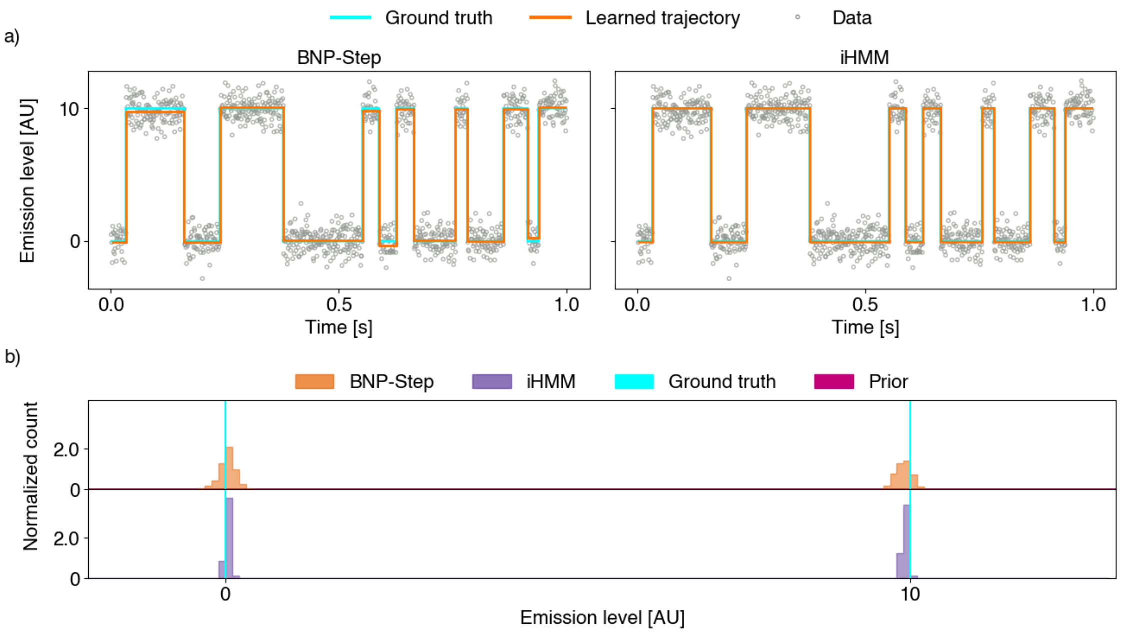


Figure S2: **Representative trajectories for two-state synthetic data from the iHMM and BNP-Step, SNR = 5.0, with conditional joint posterior distributions of emission levels for both methods.** a) Representative learned trajectories for BNP-Step and the iHMM. The same labeling and color scheme is used as in Figure S1. For BNP-Step, the orange line represents the maximum *a posteriori* (MAP) estimate. b) Conditional joint posterior distributions (as described in Supplemental Section 2) of emission levels for BNP-Step and the iHMM. For BNP-Step, the top 1000 MAP samples were used in analysis. For the iHMM, all samples after burn-in with the mode number of states were used. In this case, BNP-Step under-performs compared to the iHMM, as it incorrectly attributes fluctuations in the emission level to differences in the underlying state itself rather than to noise. This is evidenced by broader peaks in BNP-Step's posterior distribution in comparison to the iHMM.

#### 3 COMPARISON TO BIC-BASED ALGORITHM: SUPPLEMENTAL SECTION

When comparing BNP-Step to the BIC-based method of Ref. (11), we define the SNR as the ratio of the fixed step height to the standard deviation of the Normal noise. Additionally, we define the data density as the number of observations per step.

#### 3.1 Supplementary Figures

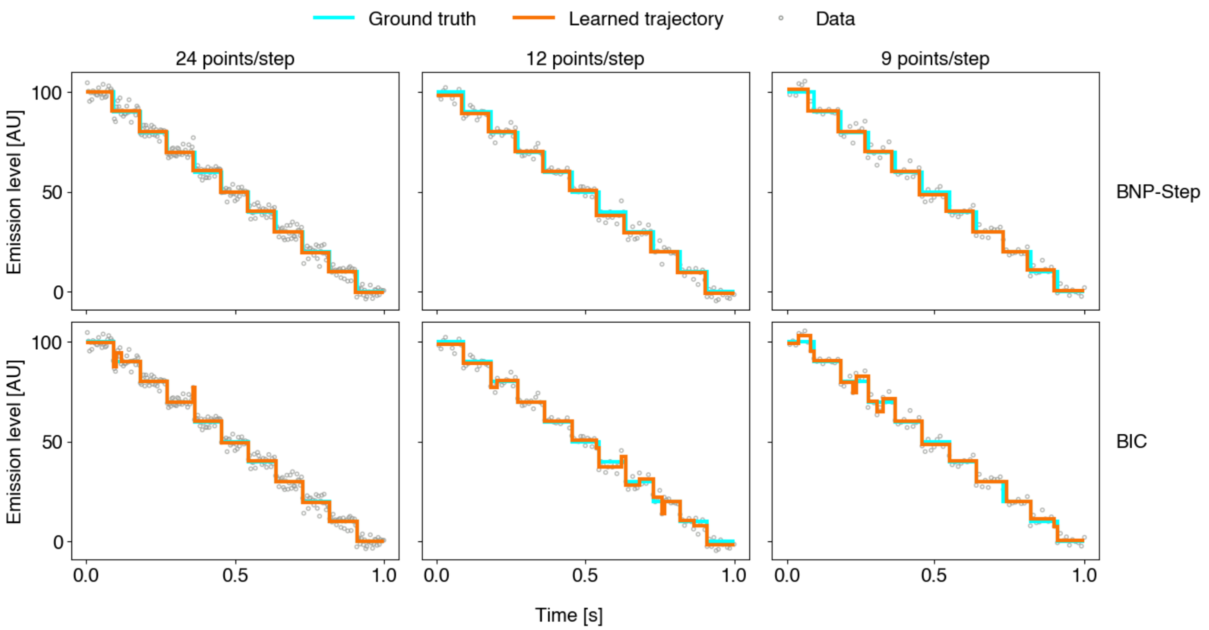


Figure S3: **Representative trajectories from BNP-Step and the KV method for synthetic data sets at data densities of 24 points/step, 12 points/step, and 9 points/step.** The top row represents BNP-Step applied to representative synthetic data sets, while the bottom row represents the KV method applied to the same sets in each column. The same color scheme is used as in Figure S1. The BNP-Step trajectories corresponds to the MAP estimates, while the KV trajectories corresponds to its point estimates. Note the increasing presence of over-fitted steps in the learned trajectories of the KV method as compared to BNP-Step as the data density decreases.

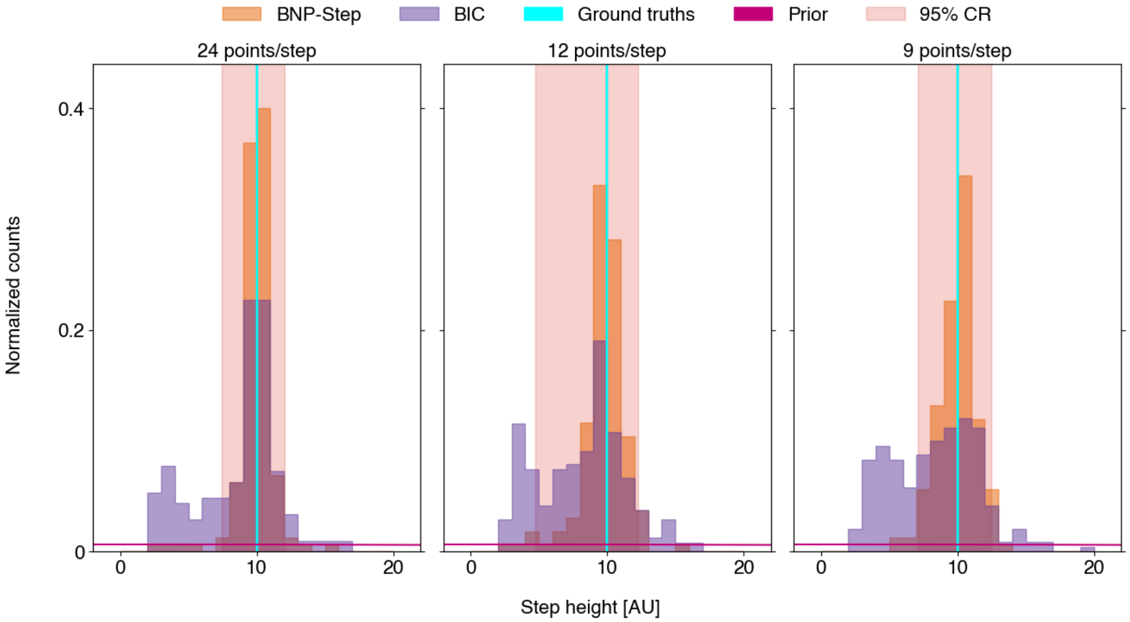


Figure S4: **Normalized histograms of sampled step heights for BNP-Step and the KV algorithm, for data densities of 24 points/step, 12 points/step, and 9 points/step.** The orange bars represent BNP-Step's MAP estimates for the analyzed synthetic data sets at each data density, while the purple bars represent the KV algorithm's point estimates for the same synthetic data sets. The cyan line represents the common ground truth value for the fixed step heights, while the magenta line represents the prior distribution used in BNP-Step.

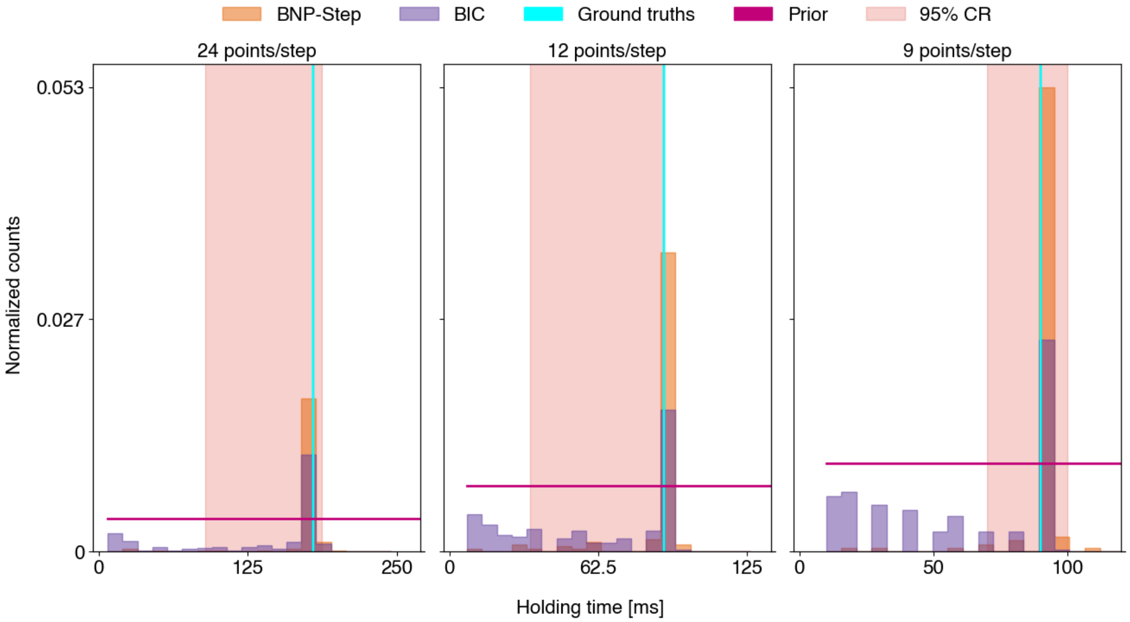


Figure S5: **Normalized histograms of sampled holding times for BNP-Step and the KV algorithm, for data densities of 24 points/step, 12 points/step, and 9 points/step.** The same labeling and color scheme are used as for Figure S4. Note the KV algorithm's shift to a bimodal distribution of step lengths as the data density decreases, with the second peak located near the inter-observation time.

### 4 APPLICATION TO EXPERIMENTAL DATA: SUPPLEMENTAL FIGURES

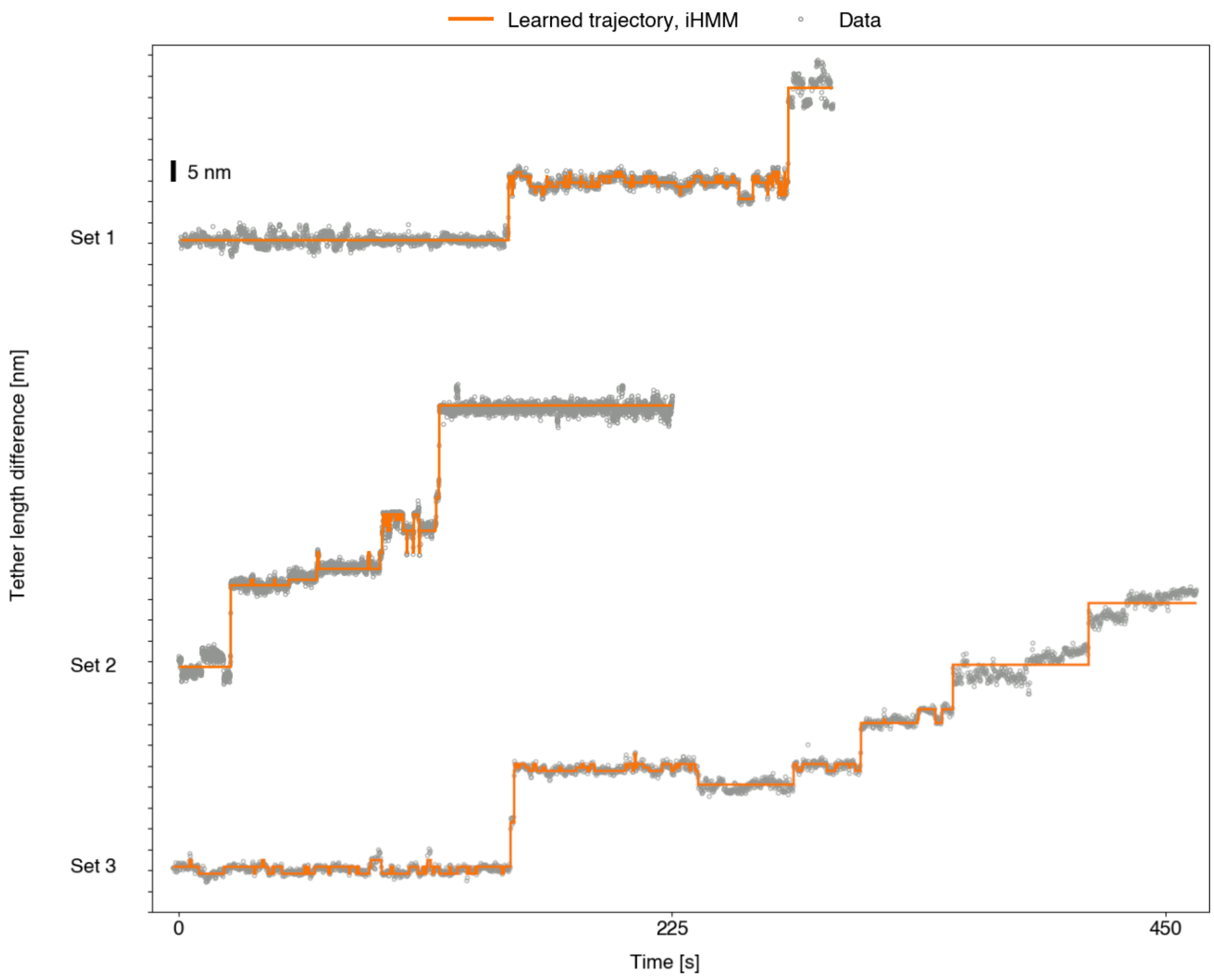


Figure S6: **Representative fits to select telomerase DNA elongation data sets for the iHMM.** The orange line represents the iHMM's mode emission mean estimate (as described in Supplemental Section 2) for each data set. While in some areas the iHMM appears to be relatively insensitive to small fluctuations in the data, in others the iHMM fits many small-amplitude fluctuations, on account of its inability to account for continuous low-frequency variations in the data.

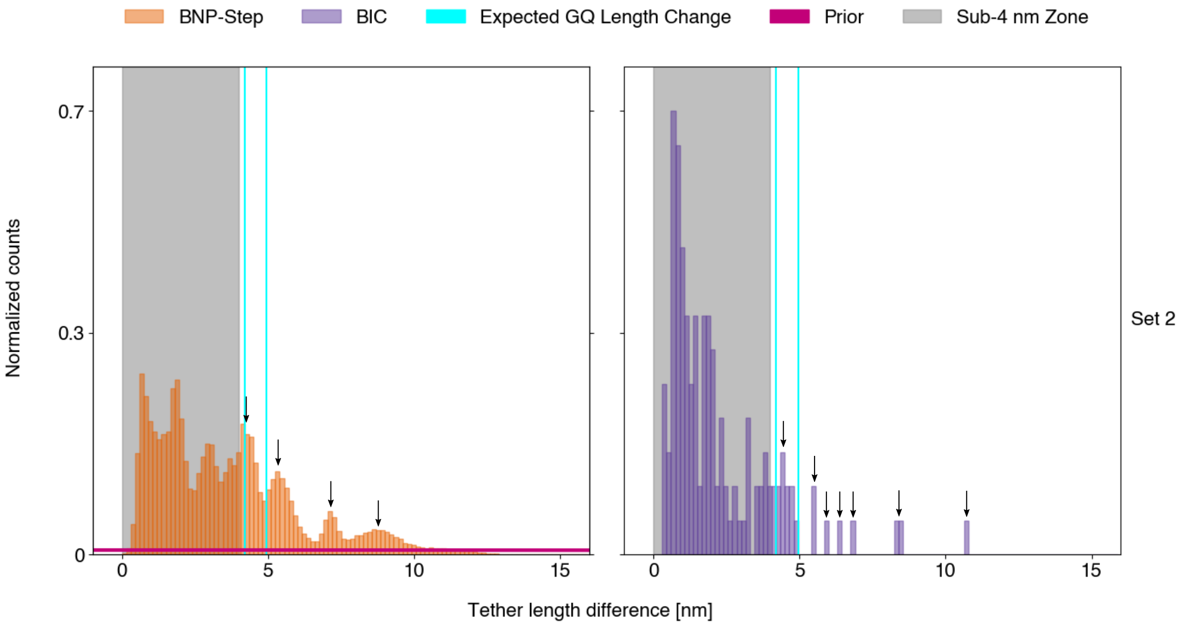


Figure S7: **Joint posterior distribution of tether length changes from BNP-Step and point estimate frequentist histogram of tether length changes from the KV method, for telomere DNA elongation data set 2 in Figure 12 of the main text.** Here, we use the same definition of the joint posterior distribution as given in Supplemental Section 2. The same color scheme is used as in Figure 13 in the main text. For BNP-Step, the top 100 MAP samples (conditioned on the MAP number of steps) for each data set were used. As in Figure 13 in the main text, distinct peaks are indicated by the small black arrows. While the peak near 4.2 nm is irregular—potentially indicating two or more closely spaced peaks—we have indicated it using only one black arrow as multiple peaks cannot be unambiguously resolved.

#### 5 ROBUSTNESS TESTING

To characterize BNP-Step's robustness, we tested its performance when the hyper-parameters were chosen to yield broad priors. In general, this did not meaningfully impact performance, to the extent that broad priors were always used in our analyses. While using physical reasoning to choose narrower priors may speed convergence, there is no inherent need to do so.

In contrast to all other hyper-parameters in our algorithm—which control the center and width of the prior distributions—the hyper-parameter $\gamma$ represents the prior probability that a given $b_m$ will equal 1. We evaluated the robustness of BNP-Step to extreme values of this hyper-parameter by testing performance on a synthetic data set at extremely low and extremely high values of $\gamma$, where the weak limit was set to the number of data points. Figure S8 shows the results of this analysis. At extremely low values of $\gamma$, performance is not impacted. At very high values of $\gamma$, substantial over-fitting is noted. Additionally, there is a pathological step at the end of the trajectory where, due to a lack of data, the height of the final step is not penalized and is allowed to grow arbitrarily large in magnitude. We conclude that values of $\gamma$ on the order of $M$ represent a region where BNP-Step breaks down. However, the use of high values for $\gamma$ may represent underlying physics that is incompatible with the use of step-finding algorithms in general.

Figure S9 represents the application of BNP-Step to a synthetic data set using different values for the weak limit $M$. So long as the weak limit is greater than or equal to the number of ground truth steps present in the data, BNP-Step's performance is not affected. If the weak limit is smaller than the the number of ground truth steps, BNP-Step fails to find a number of steps equal to the difference between the ground truth step number and the weak limit. This behavior is expected theoretically, as an inadequately large weak limit will restrict the number of steps BNP-Step is able to detect. As BNP-Step is not affected by large weak limits, this suggests that $M$ should be made as large as feasibly possible to avoid this behavior.

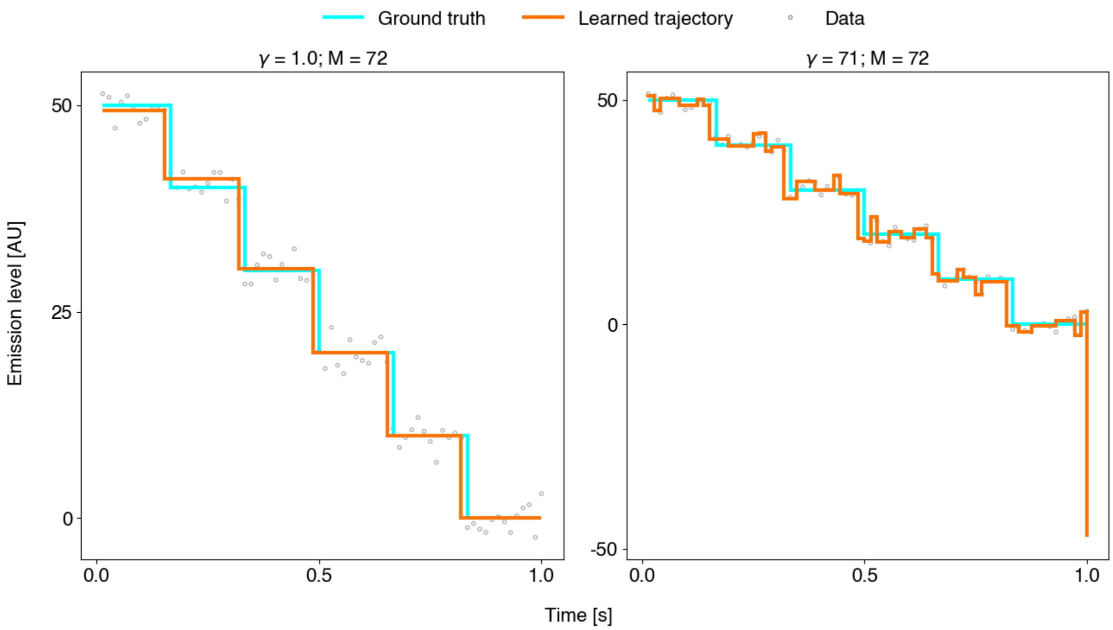


Figure S8: **Learned trajectories for 5-step synthetic data set at SNR = 4.0, for $\gamma$ = 1.0 and $\gamma$ = 71.** The same definitions and color scheme are used as in Figure S3. In each case, $M = N = 72$.

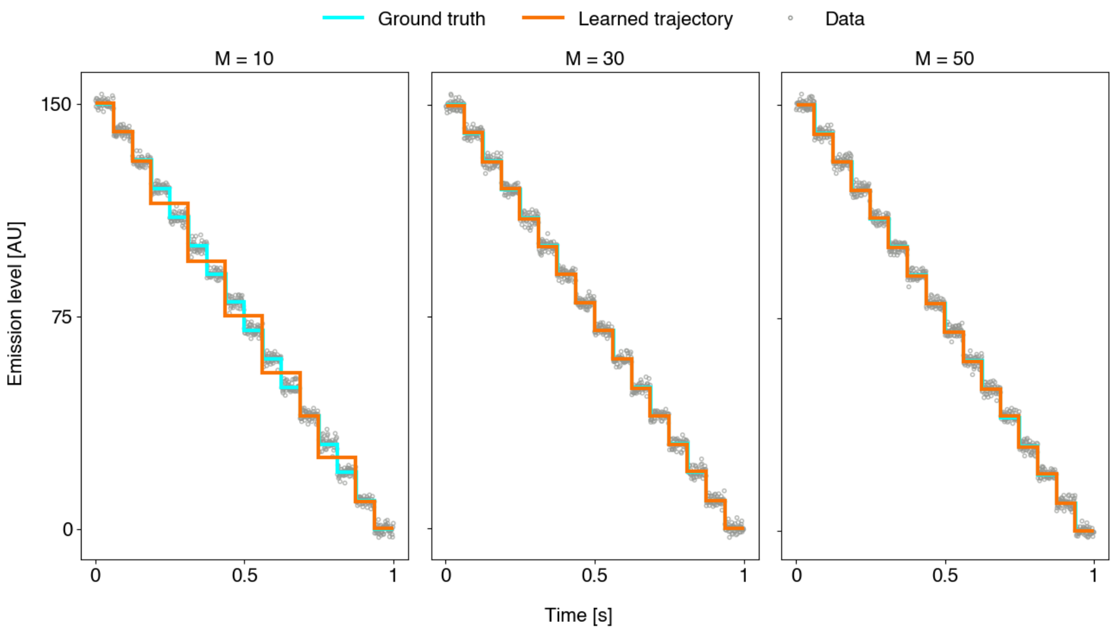


Figure S9: Learned trajectories for 15-step synthetic data set at SNR = 4.0, for $M$ = 10, $M$ = 30, and $M$ = 50. The same definitions and color scheme are used as in Figure S3.
